# Ketogenesis protects against MASLD-MASH progression through mechanisms that extend beyond overall fat oxidation rate

**DOI:** 10.1101/2024.10.17.618895

**Authors:** Eric D. Queathem, David B. Stagg, Alisa B. Nelson, Alec B. Chaves, Scott B. Crown, Kyle Fulghum, D. Andre d’Avignon, Justin R. Ryder, Patrick J. Bolan, Abdirahman Hayir, Jacob R. Gillingham, Shannon Jannatpour, Ferrol I. Rome, Ashley S. Williams, Deborah M. Muoio, Sayeed Ikramuddin, Curtis C. Hughey, Patrycja Puchalska, Peter A. Crawford

## Abstract

The progression of metabolic-dysfunction-associated steatotic liver disease (MASLD) to metabolic-dysfunction-associated steatohepatitis (MASH) involves complex alterations in both liver-autonomous and systemic metabolism that influence the liver’s balance of fat accretion and disposal. Here, we quantify the relative contribution of hepatic oxidative pathways to liver injury in MASLD-MASH. Using NMR spectroscopy, UHPLC-MS, and GC-MS, we performed stable-isotope tracing and formal flux modeling to quantify hepatic oxidative fluxes in humans across the spectrum of MASLD-MASH, and in mouse models of impaired ketogenesis. We found in humans with MASH, that liver injury correlated positively with ketogenesis and total fat oxidation, but not with turnover of the tricarboxylic acid cycle. The use of loss-of-function mouse models demonstrated that disruption of mitochondrial HMG-CoA synthase (HMGCS2), the rate-limiting step of ketogenesis, impairs overall hepatic fat oxidation and induces a MASLD-MASH-like phenotype. Disruption of mitochondrial β-hydroxybutyrate dehydrogenase (BDH1), the terminal step of ketogenesis, also impaired fat oxidation, but surprisingly did not exacerbate steatotic liver injury. Taken together, these findings suggest that quantifiable variations in overall hepatic fat oxidation may not be a primary determinant of MASLD-to-MASH progression, but rather, that maintenance of hepatic ketogenesis could serve a protective role through additional mechanisms that extend beyond quantified overall rates of fat oxidation.

## Introduction

The global prevalence of metabolic-dysfunction-associated steatotic liver disease (MASLD) is surging and is associated with mortality and multiple comorbidities.^1,2^ The natural history of MASLD is tightly linked to obesity and insulin resistance, and involves both liver-autonomous and systemic metabolic abnormalities that drive the ectopic storage of triacylglycerol (TAG) species in hepatocytes (*i.e.,* hepatic steatosis), which is strongly associated with cellular injury, inflammation, and the development of metabolic-dysfunction-associated steatohepatitis (MASH), a progressive fibrotic liver disease that markedly increases the risks of cirrhosis and hepatocellular carcinoma.^3–5^ While the severity of hepatic steatosis and the underlying metabolic disturbances precipitating TAG accretion may predict histological evidence of liver injury, steatosis alone does not account for MASLD-MASH pathogenesis. For any degree of steatosis, the development of MASH is marked by the appearance of oxidative and inflammatory stress, coupled to hepatocellular injury (*i.e.,* hepatocyte ballooning), which collectively can be quantified using the NAFLD (corresponding to the prior term referencing MASLD, nonalcoholic fatty liver disease) activity score (NAS), a histological scoring system ranging 0-8, which encompasses steatosis (0-3), inflammation (0-3) and hepatocyte ballooning (0-2). MASH is defined as a NAS score ≥ 4, and typically coincides with fibrosis, though fibrosis is not included when calculating NAS.^6^

Currently there is only one FDA approved drug available for the treatment of MASLD-MASH, while several others have failed due to poor efficacy and/or toxicity, illustrating a need to better understand the underlying drivers of hepatic fat accumulation, and the mechanistic link this has to the development of liver injury and advanced liver disease.^7,8^ Though incretin mimetics and sodium-glucose cotransporter 2 (SGLT2) inhibitors show great promise for MASLD, approaches that (i) are liver-specific, indicated for patients without diabetes or body mass index (BMI) > 30, (ii) limit the requirement for lifelong high-cost pharmacotherapy, and/or (iii) limit toxicities, remain an unmet need.^9,10^ Prior studies have correlated the accumulation of liver TAGs in obesity and/or MASLD to a rise in hepatic *de novo* lipogenesis (DNL)^11–16^, a diminution in polyunsaturated fatty acids (PUFAs)^17–21^, and an acceleration in hepatic oxidative fluxes^22–25^. Other studies have shown no changes in hepatic fat oxidation^26–29^, or have reported impairments in oxidative metabolism.^30–34^ Therefore, the lack of consensus regarding pathological alterations in hepatic oxidative fluxes throughout the natural history of MASLD-MASH progression illustrates one of the key obstacles in developing effective liver-targeted therapeutics.

Oxidative metabolism in the liver yields reducing equivalents (RE’s) in the forms of reduced nicotinamide adenine dinucleotide (NADH) and flavin adenine dinucleotide (FADH_2_), which are primarily sourced from hepatic fat oxidation via (1) the β-oxidation pathway, which generates acetyl-CoA, and (2) terminal oxidation of acetyl-CoA in the tricarboxylic acid (TCA) cycle.^35^ RE production is also directly coupled to the production of ATP, which fuels phosphoenolpyruvate (PEP)-derived gluconeogenesis (GNG) via the non-oxidative entry (anaplerosis) and exit (cataplerosis) of intermediates through the TCA cycle^36^. In both human and rodent models of MASLD, hepatic TCA cycle turnover and PEP-derived GNG are increased relative to controls,^25,36,37^ and very low density lipoprotein (VLDL) secretion increases with worsening steatosis^38,39^. The stimulation of fat disposal pathways may offload excess TAGs, however, the rise in lipid disposal is insufficient to compensate for the rate of lipid appearance, leading to ectopic accumulation of fat, which predisposes the liver to injury. In addition to VLDL secretion and terminal oxidation in the TCA cycle, ketogenesis, which produces D-beta-hydroxybutyrate (D-βOHB) and acetoacetate (AcAc), is a major conduit supporting fat disposal in the liver. Congenital deficiency of the fate-committing enzyme of ketogenesis, 3-hydroxy-3-methylglutaryl-CoA (HMG-CoA) synthase 2 (HMGCS2), is linked to hepatomegaly and fatty liver.^40,41^ Moreover, programmed ketogenic insufficiency in mice is associated with more aggressive MASLD progression.^42–46^ However, the relative rate of ketogenesis in MASLD, compared to controls, has varied among studies.^22,27,34,37,47^ As such, whether alterations in ketogenesis in human MASLD-MASH play driver, bystander, or compensating roles is debated, as are the mechanisms that link ketogenesis to oxidative metabolism and liver injury. To date, *in vivo* hepatic oxidative fluxes have only been quantified in humans with uncomplicated MASLD but have not been measured in humans with histologically confirmed MASH. To understand the metabolic drivers and/or predictors of MASLD-MASH development, progression, and resolution, we quantified hepatic oxidative fluxes in histologically confirmed patients with MASH using nuclear magnetic resonance (NMR) spectroscopy, ultra-high performance liquid chromatography coupled to mass spectrometry (UHPLC-MS), and sophisticated ^2^H/^13^C stable isotope tracing. Ketogenesis insufficient mouse models were used to determine the relationships among ketogenesis, the rate of overall fat oxidation, and liver injury. Together, our observations are consistent with the notion that while ketogenesis and fat oxidation are key inputs into the liver’s balance of lipid accretion and disposal, the driver role ketogenesis in the rate of overall hepatic fat oxidation may not be the only determinant of MASLD-to-MASH progression. Therefore, maintenance of hepatic ketogenesis could serve additional protective roles through mechanisms extending beyond its contribution to the rate of hepatic fat oxidation.

## Results

### Hepatic ketogenesis correlates with NAS in humans

During the progression of uncomplicated MASLD, reports of hepatic oxidative fluxes have been varied, with studies reporting (1) accelerations^22–25^, (2) no changes^26–29^, or (3) impairments^30–34^ in oxidative metabolism. However, oxidative fluxes have never been quantified *in vivo* in humans with histologically confirmed MASH (**Figure 1A**). To determine rates of hepatic fat oxidation and GNG in humans with MASH, participants were recruited from two clinical trials, NCT03997422 and NCT03587831. Recruited participants remained weight stable during the screening process and during the conduction of metabolic assessments. Participants with BMI ≥ 35 were screened with magnetic resonance imaging (MRI) to identify patients with a liver proton density fat fraction (PDFF) > 5%, indicative of hepatic steatosis, then a total of 16 participants were recruited to the study outlined in **Figure 1B**. After liver biopsy collection, glucose homeostasis and body composition were quantified using a frequently sampled intravenous glucose tolerance test (FSIVGTT), and dual-energy X-ray absorptiometry (DXA), respectively. Thereafter, during a 20h fast, oral ^2^H_2_O and [U-^13^C_3_]propionate tracers were delivered to enrich the circulating glucose pool across hydrogen and carbon atoms, whose positional isotopomeric labeling distribution supports modeling of relative reaction velocities (*i.e.,* flux) through hepatic intermediary metabolic pathways *in vivo.* Simultaneous to the quantification of plasma glucose ^13^C-and ^2^H-positional enrichment, the absolute rates of endogenous glucose production (EGP) (V_EGP_) and ketogenesis (V_RaβOHB_) were measured at metabolic and isotopic steady state, after a 2h infusion of [3,4-^13^C_2_]glucose and D-[U-^13^C_4_]βOHB, respectively, thereby allowing the absolute rates of hepatic intermediary metabolic pathways to be quantified **(Supplemental Figure 1)**.^35,48–50^ During the screening phase, and prior to metabolic flux assessments, a liver biopsy was collected and used to histologically grade liver health, which was collectively summarized by the NAS score, which ranged from 1-8 across all participants. All 16 patients exhibited histological signs of liver steatosis, and 14 out of 16 participants displayed signs of either hepatocyte cell injury (*i.e.,* ballooning) and/or lobular inflammation (**Figure 1C**). Ten of the 16 participants exhibited histopathological fibrosis. As expected, NAS strongly correlated with PDFF (r = 0.68, P value < 0.01) and liver fibrosis (r = 0.79, P value < 0.001) (**Figure 1D**), but did not correlate with BMI (**Figure 1E**). NAS also correlated positively with the acute insulin response to glucose (AIRg) (r = 0.68, P value = 0.005) during the FSIVGTT, suggestive of a correlation between insulin resistance and NAS (**Figure 1F**), however, NAS did not correlate with the homeostatic model assessment for insulin resistance (HOMA-IR) (**Figure 1G**).

**Figure 1.**
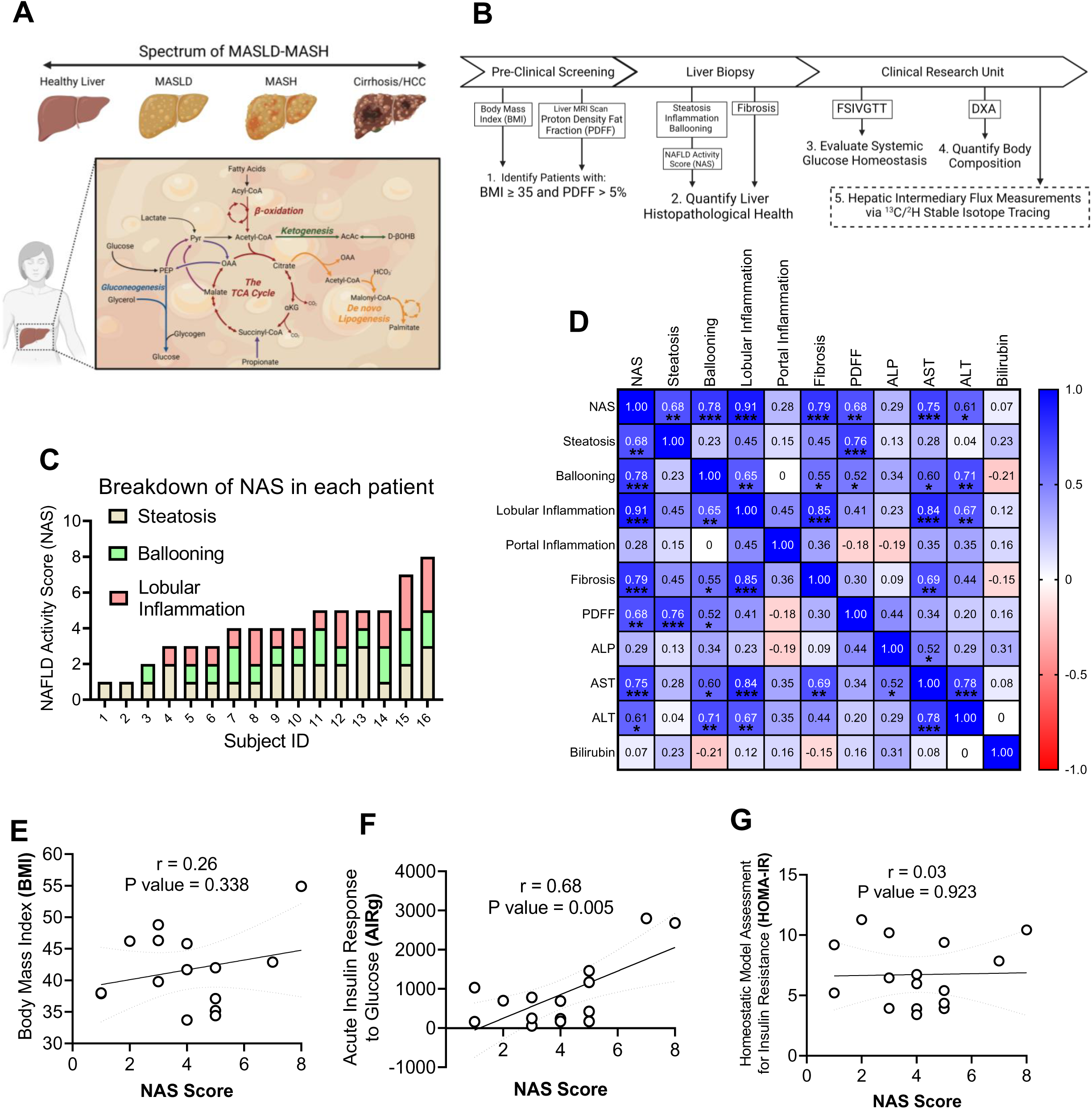
Metabolic characteristics of human participants with MASLD-MASH. **(A)** The initial stages of metabolic-dysfunction associated steatotic liver disease (MASLD) begins with simple hepatic steatosis, which has been linked to an acceleration in: de novo lipogenesis (DNL), the turnover of the tricarboxylic acid (TCA) cycle, and phosphoenolpyruvate (PEP)-derived gluconeogenesis (GNG). Metabolic shifts during the progression of MASLD to metabolic-dysfunction associated steatohepatitis (MASH) remain poorly understood. **(B)** Study design. MRI-based screening of participants (BMI ≥ 35) identified patients with a liver proton density fat fraction (PDFF) greater than 5%. Each participant underwent a liver biopsy, FSIVGTT and DXA. After an overnight fast, a panel of eight hepatic intermediary metabolic fluxes were quantified at steady-state using a multiple stable-isotope tracing approach. Metabolic fluxes were correlated with clinical markers of liver and systemic health. **(C)** Distributions of the NAFLD activity scores (NAS) in all 16 participants. **(D)** Correlation matrix of NAS with individual histological scores for steatosis, ballooning, inflammation, fibrosis, PDFF, liver enzymes and bilirubin. Correlation of NAS with **(E)** body mass index (BMI), **(F)** acute insulin response to glucose (AIRg), and **(G)** the homeostatic model assessment for insulin resistance (HOMA-IR). Pearson correlations coefficients (r) are given on each graph along with a line of best fit and 95% confidence intervals calculated using linear regression. For the correlation matrix, Pearson correlations coefficients (r) are shown in heat map format with the magnitude of the correlation given by the right-hand legend and displayed in each square. Correlations were accepted as significant if p<0.05 and are shown on each graph, or in each box on the correlation matrix an indicated, * p<0.05, ** p<0.01, ***p<0.001.

We then examined the metabolic flux modeling data acquired in this cohort **(Figure 2A-B)**. Across all 16 participants, 29 ± 8% of V_EGP_ arose from glycogenolysis (V_Glycogen_), 18 ± 7% from glycerol (V_Glycerol_) and 52 ± 7% from PEP (V_PEP_) (**Figure 2C**). Next, by simultaneously quantifying the rates of (i) TCA cycle flux, V_CS_, (ii) total ketone body (TKB) production, V_RaTKB_, and (iii) GNG sourcing pathways, V_Glycerol_ and V_PEP_, the majority of hepatic fat oxidation and RE production was quantified. The majority of RE’s produced in the fasted liver originated from β-oxidation, accounting for 43 ± 14 μmol REs/min/kg-lean body mass (LBM), while RE’s derived from the TCA cycle accounted for 17 ± 6 μmol REs/min/kg-LBM, and GNG accounted for 2 ± 1 μmol REs/min/kg-LBM (**Figure 2D**). To determine relationships among modeled hepatic fluxes, we performed regression analyses and obtained Pearson correlation coefficients **(Supplemental Figure 2A)**. As expected, V_EGP_ correlated individually with V_Glycogen_, V_PEP,_ and total anaplerosis, V_PEPCK_. Moreover, expected strong correlations among V_PEPCK_ and pyruvate cycling, V_PK+ME_ (r = 0.99, P value < 0.0001), V_CS_ and V_PK+ME_ (r = 0.60, P value = 0.014), and V_CS_ and V_PEPCK_ (r = 0.65, P value = 0.006) were observed **(Supplemental Figure 2B)**. Additional correlations were observed between V_EGP_ and V_PK+ME_ (r = 0.63, P value = 0.009), between V_EGP_ and V_PEPCK_ (r = 0.70, P value = 0.003), and between V_EGP_ and V_CS_ (r = 0.57, P value = 0.020) **(Supplemental Figure 2C)**, as well as between V_PEP_ and V_PK+ME_ (r = 0.68, P value = 0.004), between V_PEP_ and V_PEPCK_ (r = 0.79, P value < 0.001), and between V_PEP_ and V_CS_ (r = 0.68, P value = 0.004) **(Supplemental Figure 2D)**. Inverse correlations were observed between the percentage of V_EGP_ attributable to glycerol (V_Glycerol_) and V_PK+ME_, V_PEPCK,_ and V_CS_ **(Supplemental Figure 2E)**. No statistically significant correlations were observed between the ketogenic flux V_RaβOHB_ and any individual flux **(Supplemental Figure 2A)**. However, as expected, total hepatic ketogenesis (V_RaTKB_) was highly correlated with the aggregate of total hepatic fat oxidation (r = 0.99, P value < 0.001), suggesting the primary driver role of ketogenesis in determining the overall rate of hepatic fat oxidation in these conditions **(Supplemental Figure 2F)**. Interestingly, total fat oxidation was not correlated with V_CS_ (r = 0.08, P value > 0.05) **(Supplemental Figure 2G)**. Taken together, these data reflect internal consistency in the modeling data, demonstrating expected relationships between TCA cycle flux and GNG, as well as among TCA cycle flux, ketogenesis, and total fat oxidation in this cohort of participants.

**Figure 2.**
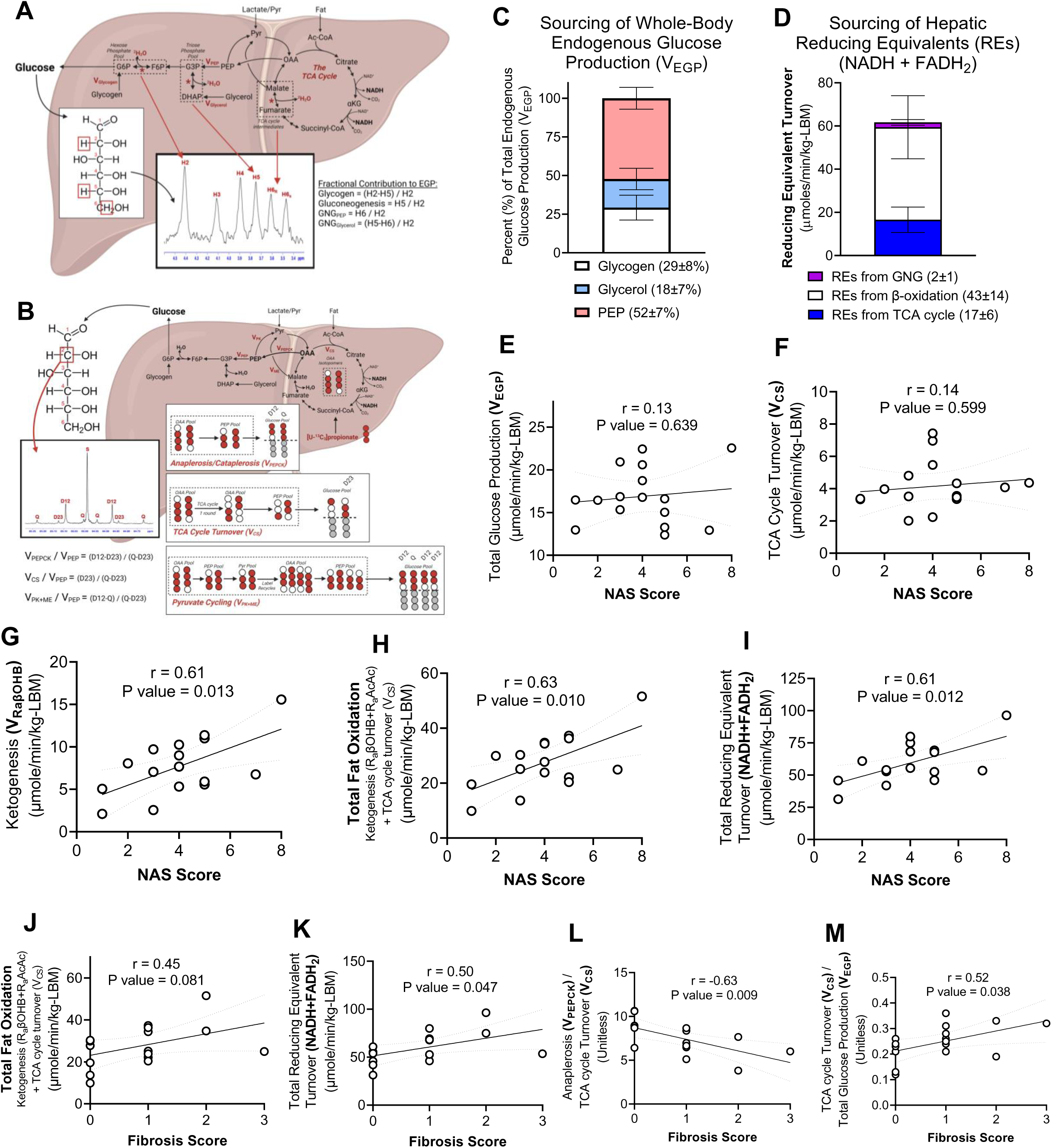
Liver injury in MASLD-MASH correlates with endogenous ketogenesis and hepatic fat oxidation. Fluxes through hepatic intermediary metabolic pathways were quantified *in vivo* non-invasively following administration of stable isotope tracers. Participants orally ingested heavy water (^2^H_2_O) and [U-^13^C_3_]propionate. [3,4-^13^C_2_]glucose and D-[U-^13^C_4_]βOHB were intravenously infused allowing whole body glucose and βOHB turnover to be quantified at metabolic and isotopic steady state. **(A)** Fractional sourcing of glucose production can be quantified from the ^2^H enrichment pattern of plasma glucose using ^2^H NMR. The product of total glucose production and fractional sourcing allow absolute reaction velocities for glucose sourcing pathways to be quantified in the liver. **(B)** Administration of [U-^13^C_3_]propionate ^13^C-enriches TCA cycle intermediates, which are in exchange with phosphoenolpyruvate (PEP). Using ^13^C NMR, and the multiplet arising from the C2 resonance of plasma glucose, the resulting metabolic network models oxidative and anaplerotic nodes of the TCA cycle in parallel to glucose production. Normalizing fluxes to the rate of PEP conversion to glucose (panel A), the absolute reaction velocities of the TCA cycle, anaplerosis, cataplerosis and pyruvate cycling can be quantified. **(C)** Average percent of endogenous glucose production (V_EGP_) derived from glycogen, glycerol and PEP across all 16 participants, highlighting that TCA cycle-sourced PEP is the major contributor to V_EGP_ in the fasted state. **(D)** Average reducing equivalents (REs) derived from GNG, β-oxidation and the TCA cycle. The correlations of NAS score with **(E)** V_EGP_, **(F)** TCA cycle turnover (V_CS_), **(G)** endogenous ketogenesis (V_RaβOHB_), **(H)** total fat oxidation, and **(I)** total RE turnover. The correlation of liver fibrosis score with **(J)** total fat oxidation, **(K)** RE production rate, **(L)** the ratio of anaplerosis relative to TCA cycle turnover (V_PEPCK_ / V_CS_), and **(M)** the ratio of TCA cycle turnover relative to total glucose production (V_CS_ / V_EGP_). Data are expressed as either mean ± standard deviation or shown as correlations. Pearson correlations coefficients (r) are shown on each group along with a line of best fit and 95% confidence intervals calculated using linear regression. Correlations were accepted as significant if p<0.05. P values are shown on each graph.

Given the validity of the flux modeling data collected, we next sought to determine the relationships among these measured hepatic fluxes and MASLD-MASH progression. Surprisingly, despite the known relationships between insulin resistance and V_EGP_ in many populations, and the relationship between insulin resistance and NAS, V_EGP_ did not correlate with NAS in this population (r = 0.13, P value = 0.639) (**Figure 2E**). Moreover, V_CS_, V_Glycogen_, V_Glycerol_, V_PEP_, V_PK+ME_, and V_PEPCK_ all showed no correlation with NAS in this population (**Figures 2F**, **Supplemental Figure 3A-B).** However, surrogates for the rate of ketogenesis correlated directly with NAS [V_RaβOHB_, **Figure 2G** (r = 0.61, P value = 0.013); and V_RaAcAc_ (r = 0.58, P value = 0.018) and V_RaTKB_ (r = 0.61, P value = 0.013), **Supplemental Figure 3C**]. Thus, consistent with the strong correlation between ketogenesis and total fat oxidation **(Supplemental Figure 2F),** both total fat oxidation (r = 0.63, P value = 0.010) and total RE production (r = 0.61, P value = 0.012) rates also correlated directly with NAS (**Figure 2H-I**). The mitochondrial redox ratio of βOHB:AcAc did not correlate with NAS **(Supplemental Figure 3D).** V_RaβOHB_ did not correlate with liver PDFF (r = 0.38, p value = 0.143), but trends towards a positive correlation with AIRg (r = 0.46, p value = 0.081) **(Supplemental Figure 3E-F**). V_RaβOHB_ also positively correlated with circulating transaminase levels [AST, r = 0.51, P value = 0.041; ALT, r = 0.52, P value = 0.037] **(Supplemental Figure 3G).** Though not included in NAS, fibrosis is also a strong prognostic indicator of MASLD progression. Total fat oxidation trended towards a positive correlation with fibrosis (r = 0.45, P value = 0.081), whereas RE production rate was significantly correlated with fibrosis (r = 0.50, P value = 0.047) (**Figure 2J-K**). Interestingly, fibrosis was inversely correlated with the rate of anaplerosis relative to the TCA cycle (V_PEPCK_ / V_CS_) (r = -0.63, P value = 0.009), and positively correlated with the rate of the TCA cycle relative to EGP (V_CS_ / V_EGP_) (r = 0.52, P value = 0.038) (**Figure 2L-M**). These ratios provide insight into the distribution of RE’s harvested from fat oxidation, and suggest that with MASLD progression, the sourcing of RE’s to glucose production may wane, consistent with the pathological course of the condition through MASH. Collectively, these data demonstrate that ketogenesis and total fat oxidation increased in this population of participants during the natural history of MASLD progression, whereas TCA cycle flux remained relatively stable. Given (i) the relationship between ketogenesis and MASLD and (ii) the fact that ketogenesis is a strong predictor of total fat oxidation in the liver, we next sought to determine the mechanistic connections among these indices using mouse models.

### Loss of HMGCS2 predisposes the liver to steatosis and dysregulated hepatic energy metabolism

Our previous findings in *ex vivo* perfused livers suggested that GNG was impaired in the setting of HMGCS2 knockdown-driven ketogenesis insufficiency.^42,49^ Because hepatic GNG is bioenergetically coupled to fat oxidation, this suggested that overall hepatic fat oxidation might be impaired in the absence of ketogenesis. Nonetheless, ketogenesis insufficient mice maintained euglycemia when fasted, which possibly is due to compensation from glycogenolysis.^42,49,51^ To test if sustained fat oxidation was impaired in the setting of ketogenic insufficiency *in vivo,* we fed control and hepatocyte-specific HMGCS2-null mice a high-fat, carbohydrate-restricted (HFCR) diet, and hypothesized that in the absence of dietary carbohydrates, hepatic GNG would fail to support glycemia (**Figure 3A**). We first confirmed loss of HMGCS2 function in hepatocyte-specific HMGCS2 knockout (KO) mice^45,52^ by demonstrating that chow-fed HMGCS2-Liver-KO mice failed to mount a ketogenic response to fasting, but maintained a normal ratio of βOHB:AcAc (**Figure 3B-C**).^53^ When littermate control mice were placed on a HFCR diet for 1 week, 4h fasted total ketone bodies were elevated 3.3 ± 0.2-fold, as expected (Student’s t-test, P value < 0.001); however, only βOHB was increased, and not circulating AcAc, which was lower in the HFCR diet-fed mice (**Figure 3D**). As a result, the ratio of βOHB:AcAc was increased 10.7 ± 0.2-fold, indicative of an elevated mitochondrial NADH:NAD^+^ ratio (**Figure 3E**). Moreover, unexpectedly, while circulating total ketones in HFCR-fed HMGCS2-Liver-KO mice (662 ± 152 μM) trended lower than littermate controls (1071 ± 349 μM) (Student’s t-test, P value = 0.08), they were markedly elevated compared to chow-fed HMGCS2-Liver-KO animals (66 ± 5 μM) (**Figure 3B, 3F**). Compared to littermate controls maintained on HFCR diet, loss of HMGCS2 markedly decreased the ratio of βOHB:AcAc in HFCR diet fed mice, but increased the total amount of circulating L-βOHB by 3.8 ± 0.1-fold, which does not contribute to mitochondrial NADH:NAD^+^ ratio (**Figure 3G-I**). To address whether the ketonemia in HFCR-diet fed HMGCS2-Liver-KO mice was attributable to ketogenesis or to possible impairments in peripheral ketone body disposal, we quantified static abundances of ketone bodies in liver tissue and found that HFCR-fed HMGCS2-Liver-KO livers showed a 69 ± 14% diminution in total ketone bodies (Student’s t-test, P value < 0.01), and increased liver L-βOHB, which together are supportive of marked impairment of conventional ketogenesis (i.e., that which produces D-βOHB) in livers of HFCR-fed HMGCS2-Liver-KO mice (**Figure 3J-K**).

**Figure 3.**
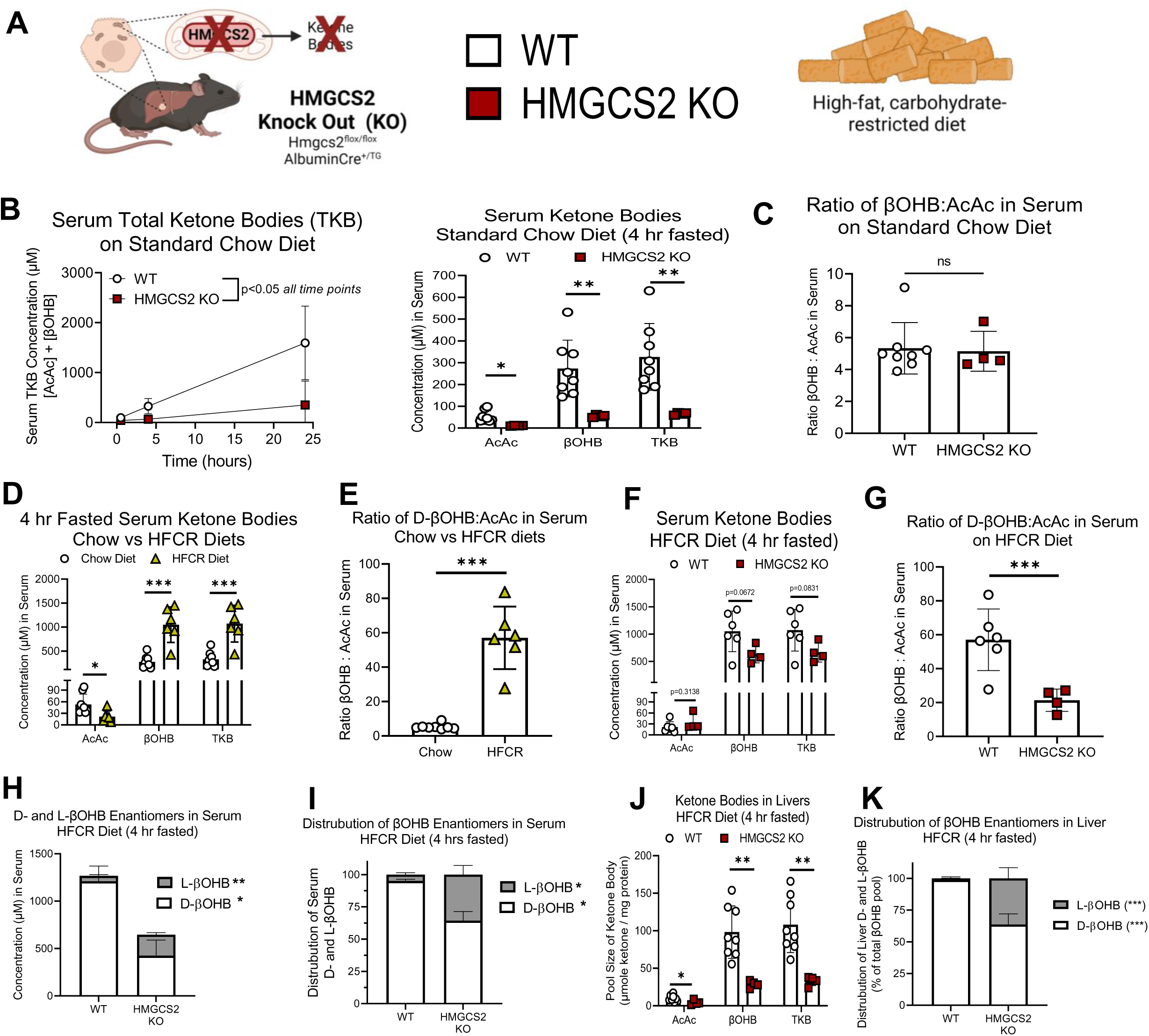
Ketogenic insufficiency induced by loss of hepatocyte HMGCS2 impairs fasting ketosis and provokes a non-physiological rise in L-βOHB in HFCR-diet-fed mice. **(A)** Ketogenesis-null mice were generated by deleting 3-hydroxymethylglutaryl-CoA synthase 2 (HMGCS2) in hepatocytes, the rate limiting step of ketone body production, by crossing homozygous *Hmgcs2*^flox/flox^ mice with transgenic heterozygous mice expressing Cre recombinase driven by the albumin promoter. Mice were maintained on standard chow diet then age-matched littermate control male wild-type (WT) and HMGCS2 knock out (HMGCS2-Liver-KO) mice were switched to a high-fat carbohydrate-restricted (HFCR) diet for ∼1 week, and studied either in the random-fed or fasted state. **(B)** Validation of functional loss of HMGCS2 was confirmed *in vivo* in standard-chow fed mice by demonstrating HMGCS2-Liver-KO mice failed to increase total ketone bodies (TKB) during a 24 hour fast (*left*), marked by a decrease in both AcAc and βOHB (*right*, 4h fasted) (n=4-8/group). **(C)** Ratio of βOHB:AcAc in 4h fasted WT and HMGCS2-Liver-KO mice fed a standard chow diet (n=4-8/group). Wild-type mice were fed a high-fat carbohydrate-restricted (HFCR) diet for one week, then fasted for 4 hours. **(D)** AcAc, βOHB and TKBs in chow or HFCR-fed mice after 4 hours fasting (n=6-8/group). **(E)** The ratio of βOHB:AcAc in chow or HFCR-fed mice after 4 hours fasting (n=6-8/group). **(F)** Serum TKB analysis in littermate control and HMGCS2-Liver-KO mice fed a HFCR-diet (n=4-6/group). **(G)** The ratio of βOHB:AcAc in littermate control and HMGCS2-Liver-KO mice fed a HFCR-diet (n=4-6/group). **(H)** Absolute and **(I)** relative enantiomeric distribution of the serum βOHB pool in littermate control and HMGCS2-Liver-KO mice fed a HFCR-diet (n=2-3/group). **(J)** Absolute and **(K)** relative TKB analysis of intrahepatic ketones from littermate control and HMGCS2-Liver-KO mice fed a HFCR-diet and fasted for 4 hours (n=4-8/group). Data are expressed as mean ± standard deviation. Statistical differences were determined by Student’s t-tests and accepted as significant if p<0.05. * p<0.05, ** p<0.01, ***p<0.001, as indicated. NS = not statistically significant.

To determine the systemic physiological response to acute HFCR-feeding of HMGCS2-Liver-KO mice, we quantified gravimetric indices and metabolite concentrations one week after transitioning from chow. HMGCS2-Liver-KO mice lost more weight than littermate controls when fed the HFCR diet, but developed severe hepatomegaly (**Figure 4A-B**). This was accompanied by a 3.5 ± 0.1-fold accumulation of liver TAGs (**Figure 4C-D**), which could be reversed by refeeding mice a standard chow-diet for one week (**Figure 4E**). In both the random-fed state, and 4 hour fasted state, HMGCS2-Liver-KO mice fed a HFCR diet for one week had elevated plasma non esterified fatty acids (NEFAs), no differences in circulating TAGs, decreased blood glucose concentrations, and markedly depleted hepatic glycogen stores (**Figure 4F-G**).

**Figure 4.**
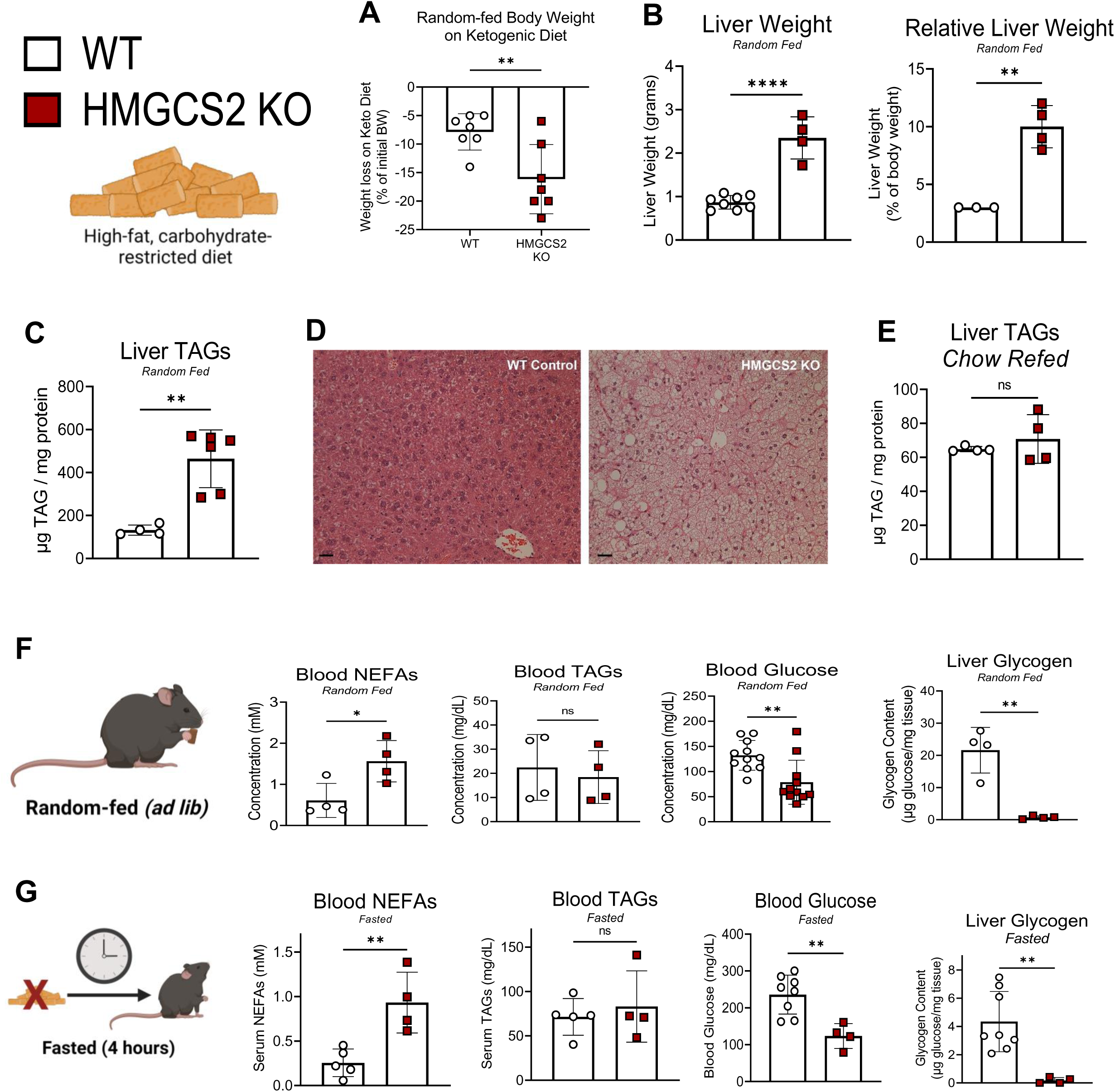
Loss of hepatocyte HMGCS2 induces hepatomegaly, liver steatosis and relative hypoglycemia in HFCR diet-fed mice. Littermate control (WT) and HMGCS2-Liver-KO mice maintained on standard chow diet were switched to high-fat carbohydrate-restricted (HFCR) diet for ∼1 week. **(A)** Net change in body weight after switching to HFCR diet (n=7/group). **(B)** Total liver weight (grams), and relative liver weight (% of body weight) (n=3-8/group). Total liver triacylglycerols (TAG) in random-fed livers quantified **(C)** colorimetrically, and shown **(D)** histologically with H&E stain. **(E)** Total liver TAGs in random-fed livers of HFCR diet fed mice, after 1 week of refeeding standard chow (n=4/group). Systemic physiological markers of whole-body metabolism in **(F)** random-fed and **(G)** 4-hour fasted mice, including: blood non-esterified fatty acids (NEFAs), TAGs, and glucose. Also included is total liver glycogen content (n=4-11/group). Data are expressed as mean ± standard deviation. Statistical differences were determined by Student’s t-tests and accepted as significant if p<0.05. * p<0.05, ** p<0.01, ***p<0.001, as indicated. NS = not statistically significant.

To test the hypothesis that the impairment of hepatic fat oxidation could be linked to hypoglycemia in HFCR-fed HMGCS2-Liver-KO mice, we quantified hepatic glucose and associated oxidative fluxes *in vivo* in conscious, unrestrained mice, using a previously established GC-MS-based approach^54–56^. To accomplish this, vascular catheters were placed in the carotid artery and jugular vein of 16-week-old mice fed a standard-chow diet. Five days into post-surgical recovery, mice were switched to HFCR diet for 2 days. Seven days post-surgery, ^2^H_2_O, [U-^13^C_3_]propionate and [6,6-^2^H_2_]glucose stable isotope tracers were intravenously infused, allowing quantification of *in vivo* fluxes via ^2^H/^13^C metabolic flux analysis (**Figure 5A**).^48,55,56^ When fasted, HFCR-fed HMGCS2-Liver-KO mice had lower rates of whole-body glucose production (V_EGP_) [59.6 ± 8.2 μmol/min/kg body weight (BW)], compared to littermate control mice (90.1 ± 12 μmol/min/kg BW, Student’s t-test, P value = 0.003, **Figure 5B**). Body weight was not different between groups (data not shown). Both littermate controls and HMGCS2-Liver-KO mice showed very low rates of glycogenolysis (V_PYGL_), however, as expected, HFCR-fed HMGCS2-Liver-KO mice showed a 31 ± 8% (Student’s t-test, P value < 0.01) diminution in total gluconeogenesis (V_Aldo_), which was linked to a reduction in sourcing of glucose from both glycerol (V_GK_) and PEP (V_Enol_) (**Figure 5C**). Gluconeogenic fluxes are tightly coupled to the entry (anaplerosis) and exit (cataplerosis) of nutrients through the TCA cycle, thus we next examined glucose linked hepatic oxidative fluxes. While anaplerosis from propionyl-CoA (V_PCC_) was decreased in HMGCS2-Liver-KO mice, most anaplerotic and associated fluxes were not different between genotypes (V_PC_, V_LDH_, and V_PK+ME_). Cataplerosis (V_PCK_) was also similar between HMGCS2-Liver-KO and WT littermates (Student’s t-test, P value > 0.05) (**Figure 5D**). However, HMGCS2-Liver-KO mice showed increased TCA cycle fluxes as evidenced by higher V_CS_ (+72 ± 6%, Student’s t-test, P value < 0.001) and V_SDH_ (+42 ± 6%, Student’s t-test, P value < 0.01) (**Figure 5D**). Thus, the V_PCK_/V_CS_ ratio was decreased 50 ± 10% (Student’s t-test, P value < 0.001) in HMGCS2-Liver-KO mice, and V_CS_/V_EGP_ ratio was increased 158 ± 8% (Student’s t-test, P value < 0.001) (**Figure 5E**). Moreover, while V_PC_ was unchanged, static concentrations of acetyl-CoA (quantified by LC-MS/MS) were increased 77 ± 17% (Student’s t-test, P value < 0.01) in livers of HFCR-fed HMGCS2-Liver-KO mice, indicating an absence of allosteric activation of PC activity due to increased acetyl-CoA in HFCR-fed HMGCS2-Liver-KO mice (**Figure 5F**). Moreover, static levels of liver succinyl-CoA and propionyl-CoA were decreased, also suggestive of diminished relative anaplerosis (**Figure 5E-F**). The total pools of energy adenylates (ATP, ADP, and AMP) were diminished 39 ± 14% in livers of HFCR-fed HMGCS2-Liver-KO mice, with no impairment in calculated energy charge **(Supplemental Figure 4A-C)**. Static liver concentrations of redox metabolites NAD^+^ and NADH were also both diminished 45 ± 13% and 38 ± 14%, respectively, but no difference in static NADH:NAD+ ratio was observed in livers of HFCR-fed HMGCS2-Liver-KO mice **(Supplemental Figure 4D-F).** Together, these findings underscore interconnections among ketogenesis, overall fat oxidation, anaplerosis/cataplerosis, and total EGP. Relative responses of modeled fluxes in HFCR-fed HMGCS2-Liver-KO mice, quantified *in vivo* via intravenous infusions, are consistent with our prior observations performed using *ex vivo* portal vein liver perfusions in chow-fed HMGCS2 knockdown mice, which revealed: (i) increased mitochondrial acetyl-CoA content, (ii) elevated V_CS_ absolute flux rate, and (iii) diminished V_PEPCK_/V_CS_ ratio.^42,49^ We next sought to quantify the contribution of ketogenesis to global hepatic fat oxidation, and how this influences the course of MASLD pathology.

**Figure 5.**
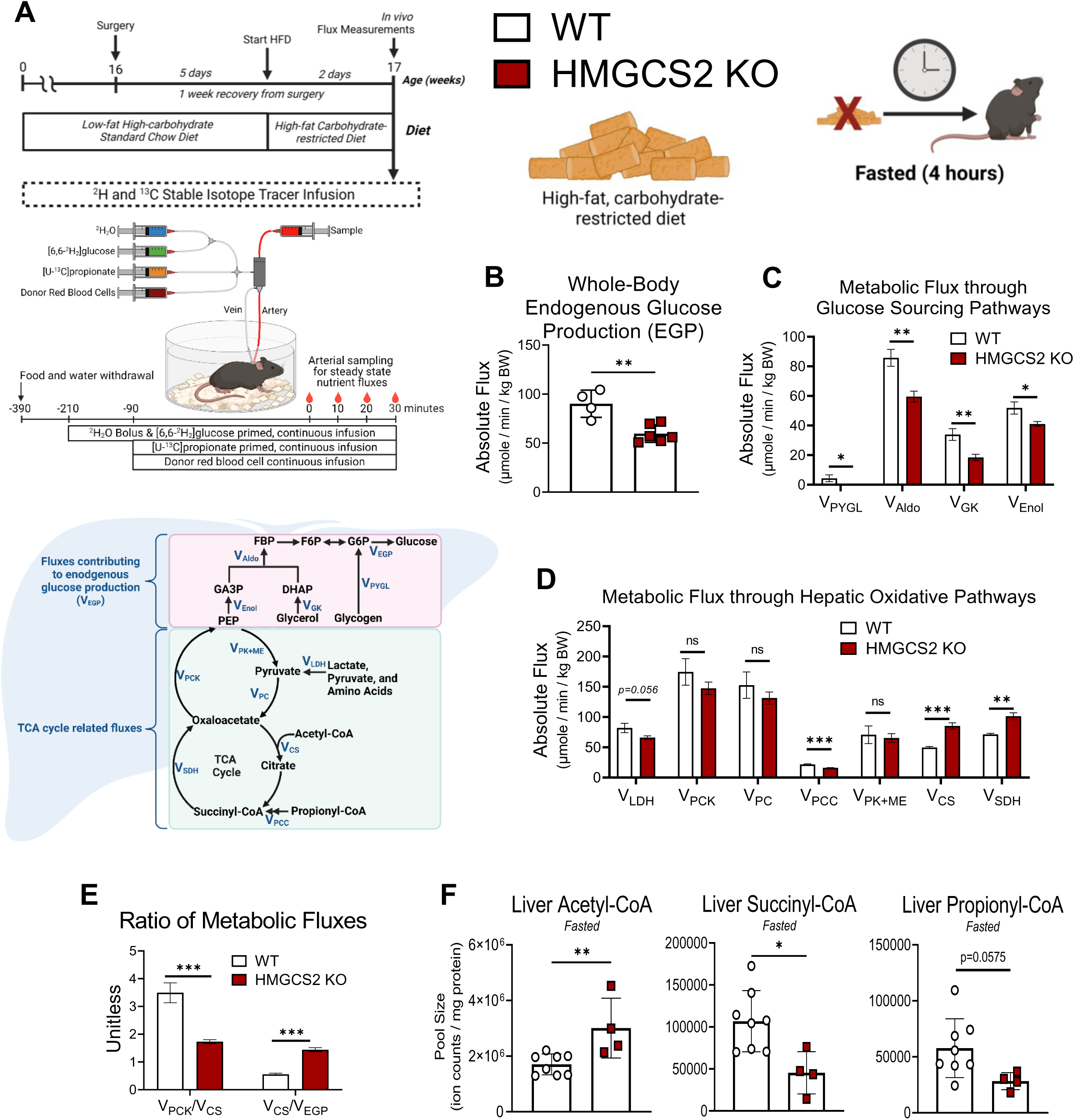
Ketogenic insufficiency diminishes gluconeogenesis in HFCR diet-fed mice, and disrupts hepatic energy metabolism in vivo. **(A)** Overview of *in vivo* hepatic flux modeling study design. *(Top)* Fluxes were measured in ∼16-week-old female littermate control (WT) and HMGCS2-Liver-KO mice. Five days after surgical implantation of indwelling cannulas into the carotid artery and jugular vein, mice were switched from standard chow to a HFCR diet, then fluxes were measured after a two-day acclimation period. *(Middle)* Multiple stable isotope tracers were administered including [6,6-^2^H_2_] glucose, [U-^13^C_3_] propionate, and heavy water (^2^H_2_O). Red blood cells from donor mice were also infused to maintain blood volume. Samples collected for flux analysis are during the final 30 minutes after metabolic and isotopic steady-state has been reached. *(Bottom)* Diagram of reaction velocities (V) (i.e., *fluxes*) modeled using a mass isotopomer distribution analysis of derivatized plasma glucose acquired via GC-MS. **(B)** Rates of whole-body endogenous glucose production (V_EGP_) at metabolic-and isotopic-steady state measured via a tracer dilution of [6,6-^2^H_2_] glucose. >90% of whole-body EGP comes from the liver, thus V_EGP_ ∼ hepatic glucose production. (n=4-6/group). **(C)** Absolute rates of glucose sourcing pathways including: glycogenolysis (V_PYGL_), total gluconeogenesis (V_Aldo_), and rates of glycerol flux to glucose (V_GK_) and phosphoenolpyruvate (PEP) flux to glucose (V_Enol_, n=4-6/group). **(D)** Absolute rates for key hepatic reactions in the oxidative flux network shown in panel A including: lactate dehydrogenase (V_LDH_), PEP carboxykinase (PCK) (V_PCK_, i.e., total TCA cycle cataplerosis), anaplerosis into the TCA cycle via pyruvate carboxylase (PC, V_PC_), anaplerosis into the TCA cycle via propionyl-CoA carboxylase (PCC, V_PCC_), pyruvate cycling as the sum of pyruvate kinase (PK) and malic enzyme (ME) (V_PK+ME_), citrate synthase flux (V_CS_, i.e., TCA cycle turnover) and succinate dehydrogenase flux (V_SDH_, n=4-6/group). **(E)** The ratio of total anaplerosis relative to TCA cycle turnover (V_PCK_ / V_CS_), and the ratio of TCA cycle turnover relative to total glucose production (V_CS_ / V_EGP_) (n=4-6/group). The pool size of various molecules that are central to hepatic energy metabolism were quantified in livers of HFCR fed fasted mice including **(F)** acetyl-CoA, succinyl-CoA and propionyl-CoA. Data are expressed as mean ± standard deviation. Statistical differences were determined by Student’s t-tests and accepted as significant if p<0.05. * p<0.05, ** p<0.01, ***p<0.001, as indicated. NS = not statistically significant.

### Ketogenic insufficiency impairs total hepatic fat oxidation

To determine if ketogenic insufficiency impaired global rates of hepatic fat oxidation (**Figure 6A**), including those components accounted for by ketogenesis and V_CS_, we returned to ketogenesis insufficient models in *ex vivo* perfused livers in carbohydrate-replete settings to avoid counterregulatory responses to hypoglycemia. Our group has previously extensively characterized the HMGCS2 knockdown model of ketogenic insufficiency via antisense oligonucleotide (ASO), and showed (i) nearly complete abrogation of HMGCS2 protein in the liver, but its preservation in non-hepatic tissues; (ii) abrogation of fasting-induced ketogenesis; (iii) elevation of V_CS_ and increase in V_CS_/V_GNG_ ratio; and (iv) increased predisposition to steatotic liver injury compared to scrambled ASO-treated controls.^42,43,49,53^ Compared to livers of scrambled ASO-treated controls, the livers of chow-fed ketogenesis insufficient mice exhibited an 80 ± 24% diminution in total fat oxidation (Student’s t-test, P value < 0.001), a 76 ± 24% impairment in REs generated from β-oxidation, and a strong trend towards a decrease in total RE production rate (39 ± 23% Student’s t-test, P value = 0.056), despite a trend toward a 70 ± 23% relative increase in TCA cycle-sourced REs (Student’s t-test, P value = 0.1108) (**Figure 6B-C**). There were no differences in REs generated through GNG. Concordant observations were observed in livers of HMGCS2 knockdown mice fed a sucrose-enriched, 42% kcal from fat Western diet for 8 weeks, in which V_CS_ was increased 155 ± 30% (Student’s t-test, P value = 0.03), and as such REs produced by the TCA cycle strongly trended towards a >2-fold increase in the HMGCS2 knockdown liver (Student’s t-test, P value = 0.05), suggesting that loss of ketogenesis stimulates terminal oxidation on Western diet (**Figure 6D-F**). Overall, however, the data support the hypothesis that the rate of total fat oxidation is impaired by ketogenic insufficiency.

**Figure 6.**
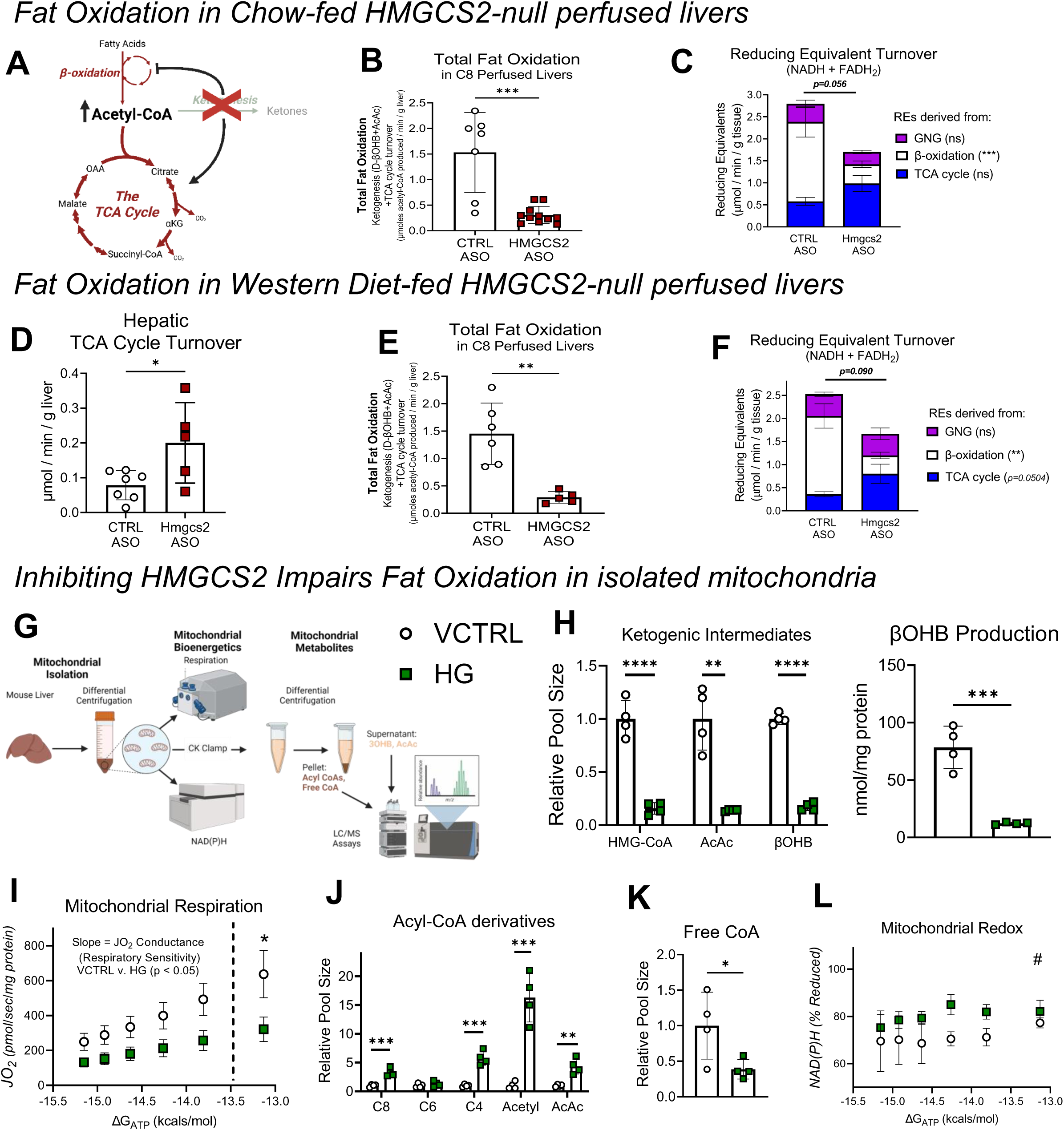
Ketogenic insufficiency impairs hepatic fat oxidation. **(A-C)** Prior studies have demonstrated that ketogenesis-insufficiency induced by loss of HMGCS2 predisposes the liver to high fat diet-induced injury, coupled with an accumulation of mitochondrial acetyl-CoA and acceleration of the TCA cycle. Livers of scrambled control antisense oligonucleotide (ASO) or mouse *Hmgcs2* ASO were perfused *ex vivo* with Krebs-Henseleit buffer supplemented with 0.2 mM sodium octanoate, 1.5 mM sodium lactate, 0.15 mM sodium pyruvate, 0.25 mM glycerol, 0.2 mM [U-^13^C_3_] propionate and 3% ^2^H_2_O, then oxidative fluxes were quantified using stable isotope tracing and formal flux modeling. First fat oxidation was studied in livers of chow fed mice. **(B)** Total fat oxidation quantified as the summation of (2 x ketogenesis) + TCA cycle turnover in perfused livers from control and Hmgcs2 ASO mice maintained on a standard chow diet (n=10-11/group). **(C)** Reducing equivalent (RE) turnover (NADH + FADH_2_) broken down into RE’s generated from gluconeogenesis (GNG), β-oxidation, and the TCA cycle, in perfused livers from control and Hmgcs2 ASO mice maintained on a standard chow diet (n=10-11/group). **(D-F)** Fat oxidation was studied in ex vivo perfused livers of ASO-treated mice maintained on a 40% high-fat high-sucrose Western diet (WD) for 8 weeks. **(D)** TCA cycle turnover in perfused livers from control ASO and *Hmgcs2* ASO mice maintained on WD (n=5-6/group). **(E)** Total fat oxidation quantified as the summation of (2 x ketogenesis) + TCA cycle turnover in perfused livers from control and Hmgcs2 ASO mice maintained WD (n=5-6/group). **(F)** RE production rate (NADH + FADH_2_) broken down into RE’s generated from GNG, β-oxidation, and the TCA cycle, in perfused livers from control and Hmgcs2 ASO mice maintained on WD (n=5-6/group). **(G)** Mitochondrial Diagnostics Workflow. Liver mitochondria isolated by differential centrifugation and fueled by 50 uM palmitoylcarnitine+5 mM α-ketoglutarate, were used to assess the effects of Hymeglusin (HG) versus its vehicle control (VCTRL) on βOHB production, respiratory kinetics, and redox potential during a creatine kinase (CK) energetic clamp. Parallel experiments were performed to obtain post-CK clamp mitochondrial pellets and supernatant for LC/MS metabolite profiling. **(H)** Relative pool size of 3-hydroxymethylglutaryl-CoA (HMG-CoA), AcAc and βOHB in mitochondria, and βOHB production measured in supernatant collected after the CK clamp bioenergetics assays (n=4/group). **(I)** Respiration (JO_2_) plotted as a function of energy demand [ΔG_ATP_ (kcal/mol), n=4/group]. Respiratory sensitivity, or JO_2_ conductance, was calculated by measuring the change in JO_2_ against the change in energy demand (ΔG_ATP_). Results of Student’s t-test of slopes between VCTRL and HG is shown on the graph. **(J)** Relative pool size of short chain acyl-CoA species, and **(K)** free coenzyme A in mitochondria (n=4/group). **(L)** Redox potential ((NAD(P)H % reduction) plotted as a function of energy demand [ΔG_ATP_ (kcals/mol), n=4/group]. Relative pool sizes shown are from mitochondria after incubation at a fixed energy demand (ΔG_ATP_ = -14.26 kcals/mol). Data are expressed as mean ± standard deviation. Statistical differences were determined by Student’s t-tests or two-way ANOVA and accepted as significant if p<0.05. * p<0.05, ** p<0.01, ***p<0.001, as indicated. # = significant by 2-way ANOVA (p <0.05). NS = not statistically significant.

To address underlying mechanisms for impaired total fat oxidation in the setting of ketogenic impairment, we isolated mitochondria from control C57BL/6NJ mouse livers and performed mitochondrial stress testing using our standard creatine kinase clamp protocol, allowing quantification of respiratory flux (*J*O_2_) across variations in energy demand (ΔG_ATP_) (**Figure 6G**).^57^ Treatment of mitochondria with hymeglusin (HG), a small molecule inhibitor of both HMGCS isoforms (HMGCS1 is cytoplasmic enzyme and is thus not included in our mitochondrial preparations) showed the expected diminution in the pool size of HMG-CoA (the product of the HMGCS2 reaction), as well as AcAc and βOHB within mitochondria, coinciding with abrogated βOHB production rates in mitochondria stimulated with palmitoyl-L-carnitine + α-ketoglutarate (**Figure 6H**).^58,59^ With increasing energy demand (decreasing absolute value of ΔG_ATP_), *J*O_2_ rates increased with a diminished slope in HG-treated mitochondria, indicating an overall respiratory impairment in the setting of ketogenic insufficiency (**Figure 6I**). This coincided with an accumulation of acetyl-CoA, AcAc-CoA, as well as C4-C8 acyl-CoA species, and a depletion in free CoA (**Figure 6J-K**). Finally, mitochondrial redox was more reduced [NAD(P)H, % reduced] across levels of energy demand in HG-treated mitochondria (**Figure 6L**). While this result may appear to contradict the normal redox ratio observed in livers of HFCR-fed HMGCS2-Liver-KO mice **(Figure S4F)**, static concentrations of total redox nucleotides may not reflect variations in cycling enzymatically-bound pools.^50,60^ Taken together, these findings show that programmed ketogenic insufficiency causes impaired total fat oxidation *in vivo*, *ex vivo*, and in isolated mitochondria. A potential contributor to this impairment includes free CoASH trapping, which impedes procession though β-oxidation and the TCA cycle, which is supported by our prior findings in the setting of ketogenic insufficiency.^42^ A second contributor to diminished fat oxidation rates is reduced mitochondrial matrix redox potential in the setting of ketogenic insufficiency, captured in this flux-based assay that quantifies redox nucleotide turnover among actively cycling enzymatic pools.

### Loss of mitochondrial βOHB dehydrogenase (BDH1) impairs hepatic fat oxidation but does not provoke liver injury

Ketogenic insufficiency via HMGCS2 loss of function causes steatotic liver injury.^42–46^ Our novel findings here show that HMGCS2 loss of function diminishes rates of global liver fat oxidation in a manner correlated with more reduced mitochondrial redox potential. To directly interrogate the role of the ketogenic reaction directly regulating mitochondrial redox potential, catalyzed by mitochondrial BDH1 (which converts AcAc and NADH to βOHB, recovering NAD^+^, in a near-equilibrium reaction) in (i) total hepatic fat oxidation and (ii) the response to steatotic liver injury, we studied mice in which BDH1 had been deleted specifically in hepatocytes (BDH1-Liver-KO mice) (**Figure 7A**). Our prior studies demonstrated impairments of oxidative metabolism in the livers of BDH1-Liver-KO mice maintained on a standard chow diet.^50^ Here, we fed mice the 42% kcal fat Western diet for 18 weeks. As expected, and consistent with observations in BDH1-Liver-KO mice maintained on a standard chow diet, circulating βOHB concentrations and total ketone bodies were modestly diminished after an 18h fast **(Supplemental Figure 5A)**. Body weights, liver weights, circulating glucose concentrations were normal **(Supplemental Figure 5B-D).** Livers were then perfused *ex vivo* via the portal vein with a physiologically relevant mixture of long-chain fatty acids (LCFAs) conjugated to bovine serum albumin (BSA). While there was no statistically significant diminution in total ketogenesis in livers of BDH1-Liver-KO mice, all ketone bodies produced by the liver emerged as AcAc, with no βOHB production (**Figure 7B**). Total glucose production was diminished 26 ± 13% (Student’s t-test, P value = 0.0390) in livers of BDH1-Liver-KO mice, which coincided with a 48 ± 12% and 30 ± 13% decrease in the sourcing from both PEP (Student’s t-test, P value < 0.001) and glycogen (Student’s t-test, P value = 0.04) to glucose, respectively (**Figure 7C-D**). Consistent with our previous findings in livers of BDH1-Liver-KO mice maintained on a chow diet, the rates of pyruvate cycling (V_PK+ME_), anaplerosis and cataplerosis (V_PEPCK_), and TCA cycle turnover (V_CS_) were all suppressed 42 ± 16% (Student’s t-test, P value = 0.0043), 15 ± 13% (Student’s t-test, P value = 0.0005), and 39 ± 17% (Student’s t-test, P value = 0.0160), respectively, in livers of BDH1-Liver-KO mice on Western diet (**Figure 7E**). Furthermore, total fat oxidation was decreased 26 ± 12% (Student’s t-test, P value = 0.0170) and RE production rate was decreased 20 ± 10% (Student’s t-test, P value = 0.0259) in the BDH1 KO liver (**Figure 7F-G**), demonstrating that loss of the NAD^+^ regenerating step of ketogenesis alone is sufficient to impair overall fat oxidation in the liver.

**Figure 7.**
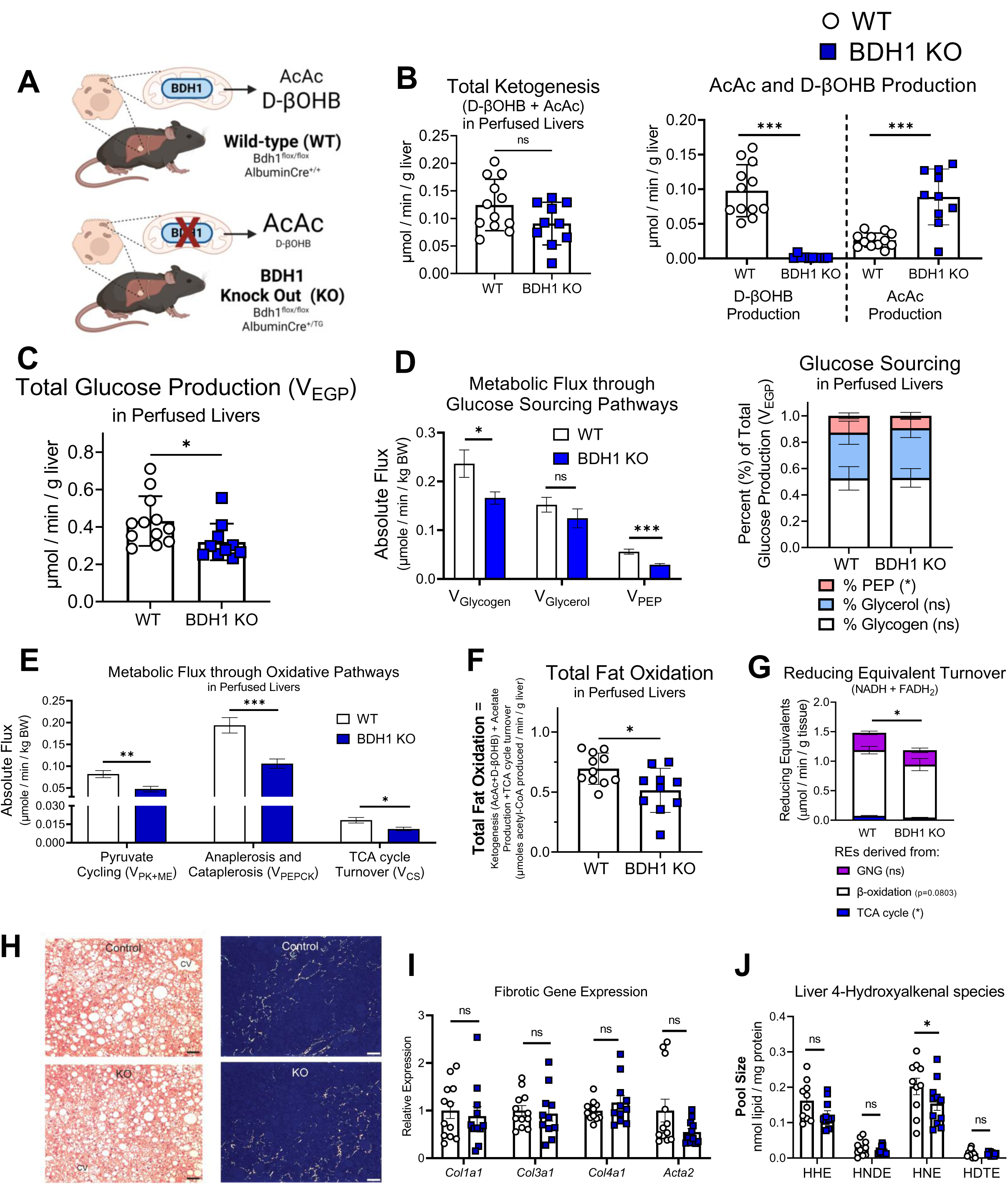
Loss of hepatocyte BDH1 impairs total hepatic fat oxidation but does not exacerbate HFD-induced liver fibrosis. **(A)** Hepatocyte-specific BDH1-null mice were generated by crossing homozygous *Bdh1*^flox/flox^ mice with transgenic heterozygous mice expressing Cre recombinase driven under the albumin promoter. Mice were maintained on standard chow diet then age-matched male littermate control (WT) and BDH-Liver-KO mice were switched to a 40% high-fat high-sucrose WD for ∼16 weeks, after which oxidation fluxes were studied in *ex vivo* perfused livers using a mixture of long-chain fatty acids (LCFAs) bound to BSA. **(B)** Total ketone body production *(left)* is normal in livers of BDH-Liver-KO mice, however, only AcAc and not βOHB is produced *(right)* (n=10-11/group). **(C)** Total hepatic glucose production (V_HGP_) and **(D)** the sourcing of glucose in LCFA perfused livers from littermate control and BDH-Liver-KO mice (n=10-11/group). **(E)** Absolute rates of oxidative pathways including: pyruvate cycling (V_PK+ME_) *(left),* total anaplerosis and cataplerosis (V_PEPCK_) *(middle)*, and TCA cycle turnover (V_CS_) *(right)* in LCFA perfused livers from WT and BDH1 KO mice (n=10-11/group). **(F)** Total fat oxidation calculated at (2 x ketogenesis) + acetogenesis + TCA cycle turnover (n=10-11/group). **(G)** Reducing equivalent (RE) turnover (NADH + FADH_2_) broken down into RE’s generated from gluconeogenesis (GNG) *(purple bar)*, β-oxidation *(white bar)*, and the TCA cycle *(blue bar)*, in LCFA perfused livers from WT and BDH1 KO mice. **(H)** Liver H&E and picrosirius histological staining of livers of littermate control and BDH-Liver-KO mice maintained on Western diet. **(I)** Relative gene expression for fibrotic gene markers measured via RT-qPCR in livers of WT and BDH1 KO mice fed a 40% WD (n=11-12/group). **(J)** Absolute levels of lipid peroxide 4-hydroxyalkenal species in livers of littermate control and BDH1-Liver-KO mice fed a 40% WD (n=10-11/group). Data are expressed as mean ± standard deviation. Statistical differences were determined by Student’s t-tests and accepted as significant is p<0.05. * p<0.05, ** p<0.01, ***p<0.001, as indicated. NS = not statistically significant.

To determine whether the diminution in hepatic fatty acid (FA) oxidation in livers of BDH1-Liver-KO mice was linked to increased hepatic steatosis or injury, we performed histological and molecular analyses. Hematoxylin and eosin staining showed normal lipid droplet storage, and no evidence of increased fibrosis via picrosirius red staining compared to littermate controls (**Figure 7H**). Gene expression biomarkers of fibrosis (*Col1a1, Col3a1, Col4a1, Acta2*) were normal, and quantities of tissue lipid peroxides (hydroxyalkenals, indices of oxidative stress) were not increased in livers of BDH1-Liver-KO mice (**Figure 7I-J**). Maintaining BDH1-Liver-KO mice and littermate controls on a 60% high fat diet (HFD), rather than a sucrose-enriched, 42% kcal fat Western diet, also showed no evidence of increased steatosis, inflammation, or fibrosis – with a trend towards a diminution in fibrosis in livers of BDH1-Liver-KO mice (56 ± 30%, Student’s t-test, P value = 0.06) **(Supplemental Figure 6A-C)**. Moreover, gene expression biomarkers of fibrosis and inflammation (*Nox, Emr1, and Tnfa*) were all diminished in livers of BDH1-Liver-KO mice **(Supplemental Figure 6C).** Collectively, these findings demonstrate (i) that homeostatic ketogenesis is required to maintain overall hepatic fat oxidation rate and its coupling to hepatic glucose production, with loss of either HMGCS2 or BDH1 showing impairments in overall hepatic fat oxidation rate, and (ii) that the impairment in overall fat oxidation rate does not fully explain why loss of HMGCS2 predisposes the liver to HFD-induced liver injury.

## Discussion

The natural history of MASLD-MASH proceeds through parallel derangements of both liver-specific and systemic metabolic pathways, as well as inflammatory and immunomodulatory signaling cascades among many cell types. Characteristic metabolic signatures include imbalanced lipid synthesis [DNL, and TAG synthesis from DNL-sourced FAs and free FAs (FFAs) derived from adipose lipolysis] and disposal (mitochondrial fat oxidation, and TAG secretion as VLDL). While most studies concur that DNL is augmented during the progression of MASLD-MASH, and VLDL secretion, while increased, does not match the rate of lipid accumulation, controversy surrounds variations in the rate of mitochondrial fat oxidation and whether variations in this rate have a causal role. This significantly impacts the selection of therapeutic approaches and molecular targets worthy of pursuit. In this study, we quantified hepatic oxidative fluxes in humans with histologically confirmed MASLD-MASH and demonstrated that the histopathological degree of liver injury (via NAS) was positively correlated with ketogenesis and total fat oxidation, but was not correlated with TCA cycle turnover. Use of mouse models suggested a driver role of ketogenesis in overall hepatic fat oxidation and a causal role in preventing liver injury, as elimination of HMGCS2 showed a decrement in overall hepatic fat oxidation. This decrement correlates with increased steatotic liver injury as shown in numerous studies performed by our group and others in the setting of ketogenic insufficiency due to deletion of HMGCS2.^42–46^ Thus, the increase in ketogenesis observed in patients with worsening MASH may serve as a compensatory mechanism, increasing fat disposal through oxidation, in the face of stagnating TCA cycle flux. This position is challenged, however, by the results observed in mice lacking BDH1 in hepatocytes, whose deficiency also quantitatively diminishes overall hepatic FA oxidation rates, but is not associated with worsened steatotic liver disease. In fact, signatures of a modestly adaptive response to diminished hepatic FA oxidation is evident in the livers of BDH1-Liver-KO mice. Taken together, these results suggest that while maintenance of ketogenic sufficiency may be important to slow the progression of MASLD, the salutary mechanism conferred by ketogenesis may not only relate to maximizing the overall rate of hepatic FA oxidation. Therefore, approaches aimed primarily at stimulating further hepatic fat oxidation may not ameliorate the course of commonly-observed MASLD, and could exacerbate liver injury.

Quantified rates of hepatic fat oxidation in humans with MASLD, relative to steatosis-free, but weight-matched controls, have been variously reported in the literature.^22–25,30–37,47,61,62^ Much of this may relate to the populations studied, including (i) the stage of progression; (ii) the degree of adiposity and insulin resistance; (iii) whether or not the participant was weight-neutral at the time of study; (iv) the genotype (*e.g., PNPLA3*); (v) whether or not the participant was fed, fasting, or undergoing a clamp study; and lastly, (vi) the specific methodology employed to quantify fat oxidation. While *in vivo* hepatic oxidative fluxes have been quantified in humans with uncomplicated MASLD, and isolated mitochondrial function has been measured in liver biospecimens harvested from participants with MASH, *in vivo* hepatic oxidative fluxes had not been quantified in humans with histologically confirmed MASH. In well-performed studies by the Burgess and Browning group, formally modeled ketogenesis negatively correlated with worsening hepatic steatosis (quantified by PDFF) in overweight and obese participants with BMI ≤ 35.^22,37^ Moreover, circulating ketones and expression levels of ketogenic enzymatic machinery decreased within worsening NAS in a bariatric surgery cohort studied by the Rector group, but no study has tested whether endogenous ketogenesis correlates with histopathologically-confirmed liver injury.^33,34^ Consistent with our findings, a recent study in non-obese participants with MASLD by the Petersen and Shulman group showed that participants with compounding cardiometabolic disturbances have increased rates of βOHB turnover in comparison to participants with simple steatosis alone, or with healthy livers, but liver biopsies were not available in that study.^27^ The cohort studied in our experiments reflects a patient population with advanced obesity at elevated risk of progression from MASH to cirrhosis, and among the metabolic fluxes modeled, the only direct correlate of NAS was endogenous ketogenesis (V_RaβOHB_). We propose that the escalation of TCA cycle flux early in the progression of MASLD may become attenuated with further progression, and indeed, the two participants we studied with the greatest V_CS_ magnitude (>5 μmol/min/kg-LBM) had NAS 4, and those exhibiting a NAS > 4 did not exceed 5 μmol/min/kg-LBM. Thus with respect to variation of TCA cycle flux, our data are generally consistent with those of the Burgess and Browning group.^22,25^ Our data in both humans and mouse models suggest that ketogenesis may serve an adaptive counterregulatory role, especially with further MASH progression.

Our findings in mouse models of altered ketogenesis suggest that impairments in rate of overall hepatic fat oxidation in ketogenic insufficiency does not relate in a direct manner to liver injury, implying that the link between ketogenesis and MASLD-MASH progression involves attributes of ketogenesis beyond its contribution to the overall rate of hepatic fat oxidation. Concordance between our observations in modeled hepatic fluxes of the human participants with MASH and mouse models of ketogenic insufficiency strongly substantiate this claim. In addition to the direct association between ketogenesis and human NAS, increasing histopathological evidence of fibrosis was inversely correlated with V_PEPCK_/V_CS_ ratio and directly with V_CS_/V_EGP_ ratio (**Figure 2L-M**), suggestive of decoupling between TCA cycle flux and sourcing of its derived REs to GNG with increasing liver injury. Similarly, V_PCK_/V_CS_ ratio was decreased in steatosis and injury-prone HMGCS2-Liver-KO mice, and V_CS_/V_EGP_ ratio was increased (**Figure 5E**). Conversely, this imbalance was not observed in livers of BDH1-Liver-KO mice, whose V_PEPCK_, V_EGP_, and V_CS_ were all coordinately diminished (**Figure 7E**). Alternatively, and not mutually exclusively, mechanisms linked to the increase in AcAc production in livers of BDH1-Liver-KO mice could explain why the loss of BDH1 protects the liver from injury.^43^

Recently, we reported a novel role for ketogenesis in supporting FA elongation, PUFA biosynthesis and phospholipid metabolism in the liver.^45^ We demonstrated that (i) ketogenesis furnishes hepatocytes with recycled acetyl-CoA that can be used for FA biosynthesis, via both the DNL and FA elongation pathways, and (ii) that incorporation of ketone body-derived carbon into FA elongation requires the cytosolic enzyme acetoacetyl-CoA synthetase (AACS), through a metabolic pathway independent of ketone body sourcing of DNL. Furthermore, we demonstrated that loss of HMGCS2 impairs FA elongation and diminishes hepatic PUFAs, specifically those esterified in phosphatidylethanolamine (PE) pools, suggesting the ketogenesis could influence mitochondrial function, and VLDL assembly and secretion through PUFA-dependent mechanisms, though these possibilities have not yet been tested. While it is tempting to speculate that the pathway linking ketogenesis to PUFA biosynthesis explains the protective nature of ketogenesis in MASLD-MASH progression, the observation that AcAc production is increased in the absence of BDH1 suggests that FA elongation and PUFA biosynthesis could be stimulated in a substrate-driven manner. PUFAs inhibit DNL in the liver via inhibiting SREBP1c mediated gene transcription,^63–68^ and therefore the rise in ketone body-sourced PUFAs in the BDH1 KO could drive both acute and chronic inhibition of DNL, while simultaneously stimulating VLDL secretion. This hypothesis is further supported by the observation that loss of HMGCS2, which abrogates production of all ketones and exhibits diminished liver PUFAs, induces a DNL gene expression profile.^42,45^

Collectively, these findings in humans with MASH and in loss-of-ketogenesis function mouse models suggest that ketogenesis may compensate in a fat-overloaded, progressively injured liver, thereby maintaining rates of overall fat oxidation, but perhaps just as importantly, by burning fat through a metabolic pathway that confers additional benefits that terminal oxidation in the TCA cycle may not bestow during MASLD progression. The specific regulatory mechanisms, whether deleterious or salutary, that coordinate the fate of β-oxidation-derived acetyl-CoA through the TCA cycle or ketogenesis, remain incompletely understood. Classical mechanisms include oxaloacetate availability, allosteric regulation of citrate synthase, NADH redox potential, and ΔG_ATP_. Physiological variation in the abundance of HMGCS2 protein could serve as a regulatory mechanism, but this has not been definitively proven. Irrespective, the pathways through which endogenous ketogenesis and exogenous ketone bodies could influence DNL, PUFA synthesis, VLDL secretion, fat oxidation, and signaling through inflammatory and immune pathways, opens opportunities to understand the range of therapeutic opportunities for metabolic and signaling roles to ameliorate MASLD.

### Limitations

The human studies are limited by a small sample size (16 participants), and the inability to study sex as a biological variable (most recruited participants were female). However, this is the first population in which isotope tracers have been used to model oxidative fluxes in participants with a BMI ≥ 35 and histopathologically graded NAS. Furthermore, while all participants were insulin resistant, only one participant had diabetes, so it is not possible to make conclusions relevant in the scope of MASLD within diabetes. Two independent experimental approaches were leveraged to delete HMGCS2 protein in mice, but the concordance in phenotypes across these models supports the rigor of our findings. Using metabolic flux modeling, we demonstrated that disruption of ketogenesis impaired overall hepatic fat oxidation, which was quantified primarily via an indirect flux modeling-based assay predicated on the assumption that mitochondrial acetyl-CoA turnover at steady-state is mathematically equivalent to total ketogenesis (V_RaTKB_) + TCA cycle turnover (V_CS_) + Acetogenesis (V_Acetate_). A limitation of this approach is the indirect measurement of fat oxidation, the assumption that fat oxidation is primarily sourced from exogenous fat sources during portal vein perfusions, and the assumption that octanoate sources little acetate. The findings generated using this methodology, however, are reinforced by the concordance among all the fluxes modeled across all orthogonal experimental methodologies employed here. Collectively, an impairment in fat oxidation was consistency observed in the setting of ketogenic insufficiency, as shown across various models including: (i) *in vivo* fluxes in HFCR-diet-fed HMGCS2-Liver-KO mice; (ii) *ex vivo* portal vein octanoate-perfused livers from HMGCS2 ASO-knockdown mice maintained on either chow or western diet; (iii) *ex vivo* portal vein octanoate-perfused livers from BDH1-Liver-KO mice maintained on chow diet; (iv) *ex vivo* portal vein LCFA-perfused livers from BDH1-Liver-KO maintained on Western diet; and (v) *ex vivo* isolated mitochondria treated with HG, which inhibits HMGCS2 and simultaneously impaired *J*O_2_ linked to fat oxidation. Collectively, these data strongly support the hypothesis that ketogenesis plays an integral role in regulating fat oxidation. A modest limitation of *in vivo* ketogenesis measurements in humans was that ketogenesis was modeled as a single pool, with V_RaAcAc_ calculated from V_RaβOHB_ and the ratio of βOHB:AcAc. Future experiments will infuse independent [^13^C]AcAc and [^13^C]βOHB tracers to directly measure V_RaAcAc_, and interconversion of AcAc and βOHB through BDH1 via a two-pool model. Finally, though male and female mice were used in these studies, as delineated in the figure legends, sex as a biological variable was not formally addressed in these studies.

## Materials and Methods

### Human MASLD-MASH recruitment and study design (Figure 1 and Supplemental Figure 1)

Patients with abnormal liver function, prediabetes or type 2 diabetes were screened to identify participants who meet eligibility requirements for two clinical studies (ClinicalTrials.gov, NCT03997422 and NCT03587831). The full list of initial inclusion and exclusion criteria are provided in **Supplemental Table 1.** Sixteen participants with BMI ≥ 35 and liver PDFF > 5%, quantified by MRI were enrolled and were not given instruction or counseling to lose weight. Insulin sensitivity and body composition were assessed via a FSIVGTT and DXA imaging, respectively, followed by *in vivo* hepatic oxidative flux measurements. After enrollment, and prior to flux measurements, a liver biopsy was obtained by an interventional radiologist under direct image guidance with ultrasound-guided approach with percutaneous puncture using a 16-gauge Tru-cut needle with standard protocols in place to detect complications. Histological severity was scored independently by two blinded pathologists based on hematoxylin and eosin-stained sections, assessing for macrovesicular steatosis (0-3), lobular inflammation (0-3), and acinar zone 3 hepatocellular injury, or ballooning degeneration (0-2), which were summed to express the NAFLD activity score, NAS. In addition to NAS, fibrosis (0-4) was also quantified using picrosirius red staining.

### PDFF Measurements

Each participant underwent a 20-minute MRI scan to measure liver fat content non-invasively. MRI scans were performed on a 3T Prisma MR system (Siemens Medical, Erlangen, Germany) at the UMN Center for Clinical Imaging Research. The PDFF image (MRI-PDFF) was acquired using a 3D GRE with 6 TE values (TR=9ms,flip angle=4, TE=1.15, 2.46, 3.69, 4.92, 6.15, 7.38 ms, 2x2x3.5mm resolution, one 17s breath-hold), and reconstructed using the vendor’s software to produce a PDFF map.

### Assessment of Glucose Homeostasis via FSIVGTT

For three days prior to admission to the clinical research unit (CRU), participants consumed a weight-maintaining diet formulated to approximate the previous 3-day dietary record. An evening meal consisting of 40% daily caloric intake and 40-50% carbohydrate and 30-40% fat was consumed before the 0700 CRU admission for FSIVGTT. Two intravenous catheters were placed, one for sampling and the other for administration of dextrose and insulin. Glucose, insulin and C-peptide were quantified at baseline and multiple times over the course of 180 minutes. HOMA-IR was calculated using the equation, HOMA-IR = fasting insulin (mU/L) × fasting glucose (mg/dL) / 405. At time 0, dextrose 0.3 grams/kg BW was given intravenously. Insulin, 0.03 units/kg BW was then injected at 20 minutes to augment glucose disposal. Insulin and C-peptide levels were measured for 10 minutes and the area under the curve was calculated as a measure of insulin secretory function, termed AIRg. A minimal model assessment was done to calculate total body insulin resistance (Si), then the Disposition Index (DI) was calculated as the product of AIRg x Si. For FSIVGTT modeling, glucose was measured via UV test enzymatic reference method with hexokinase, insulin was measured via electrochemiluminescence immunoassay (ECLIA), and C-peptide was measured via chemiluminescent immunoassay.

### Body Composition Measurements via DXA Imaging

Body composition was measured by a DXA scanner, the GE Healthcare Lunar i (GE Healthcare Lunar with enCORE software version 16.2).

### Stable isotope delivery in humans

After FSIVGTT and DXA scan, participants remained in the CRU and received a low-carbohydrate lunch meal. Participants then began fasting and did not receive an evening meal. Stable isotope intake began at 2200 hours. Participants were provided with three equal doses of 70% deuterated water (^2^H_2_O, D_2_O, 5 g/kg body water, calculated as 60% of BW in men and 50% of BW in women) every 4 hours between 2200 and 0600. These doses, along with 0.5% ^2^H_2_O available for ad libitum consumption, sufficiently labeled body water for measuring flux parameters. From 0800 to 0900 on day 2, three doses of [U-^13^C_3_]propionate (200 mg/dose) were administered orally to label TCA cycle intermediates for profiling of hepatic mitochondrial fluxes. At 0900, intravenous stable isotope infusion began with [3,4-^13^C_2_]glucose [0.563 mg/kg BW bolus, immediately followed by a two-hour infusion (0.00563 mg/min/kg BW)] and [U-^13^C_4_]βOHB [1 mg/kg BW bolus, immediately followed by two-hour infusion (0.01 mg/min/kg BW)]. At the end of the 2h infusion, ∼50 mL of blood was drawn, centrifuged to isolate serum, then aliquots were frozen at -80°C until further analysis.

### Hepatic Flux Modeling Sample Preparation and ^2^H/^13^C NMR Data Acquisition

The positional ^13^C and ^2^H isotopomers of plasma glucose encodes quantitative information about hepatic biochemical pathways, thus allowing for noninvasive modeling of hepatic metabolism *in vivo.* Direct quantification of the positional isotopomer populations of plasma glucose was achieved using ^13^C and ^2^H NMR, post derivatization into 1,2-isopropylidene glucofuranose (monoacetone glucose [MAG]) as previously described.^25,36,48,49^ MAG was reconstituted in 510 µL acetonitrile spiked with 5% ^2^H-depeleted H_2_O, then was transferred to a 5 mm 528-pp Wilmad NMR tube (Vineland, NJ) and analyzed on a Bruker Avance III 600 MHz NMR instrument equipped with a CryoProbe. During ^2^H NMR data collection, the magnet was not maintained on a field-frequency lock, as the lock channel is used to acquire ^2^H signal. Data were collected at 25°C over 1380 Hz using a 68 µs (90°) pulse, 1.3s delay between transients and Waltz16 decoupling on the proton channel during a 1s acquisition period. Depending on signal-to-noise (S/N), 4,000-40,000 transients were collected. Magnet stability was sufficient to result in deuterium line-widths of 2-3 Hz, allowing for integration to establish relative ^2^H enrichments at the number 2, 5 and 6 positions of ^2^H-labeled MAG, reflecting the relative ^2^H enrichments of plasma glucose. For ^13^C data collection, 40 µL of d_3_-acetonitrile was spiked into each sample used to collect the ^2^H NMR spectra, then a field-frequency lock was established. For data acquisition, a 12 µs excitation pulse with 22,320 Hz collection window and waltz16 decoupling for ^1^H was used. Depending on S/N, 4,000 to 40,000 transients were collected, then signal magnitudes were established by integration. The ^13^C isotopomers that appear in glucose provide a readout on oxidative metabolism in the liver, which are encoded in the multiplet pattern arising at the C2 resonance of MAG, centered at ∼85 ppm. The relative populations of the D12 doublet, D23 doublet and Q123 quartet that arise when carbons 1 and 2, carbons 2 and 3, or carbons 1, 2 and 3 are labeled, respectively, then were used to mathematically model TCA cycle turnover, anaplerosis, cataplerosis and pyruvate cycling as described below.

### Quantifying Whole-body βOHB Turnover

βOHB turnover was calculated at metabolic and isotopic steady-state from the dilution of intravenously infused D-[U-^13^C_4_]βOHB. Briefly, unlabeled and [U-^13^C_4_]βOHB were quantified using UHPLC-MS/MS, then the fractional ^13^C enrichment of plasma βOHB and tracer infusion rate were used to calculate whole-body turnover rates of βOHB using the equation:

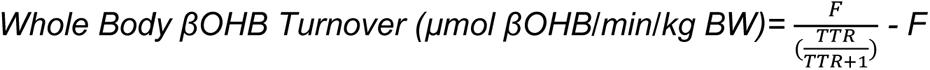

Where F is the tracer infusion rate and TTR is the Tracer : Tracee ratio ([U-^13^C_4_]βOHB / [^12^C]βOHB). After flux measurements, turnover rates are corrected for LBM from DXA imaging, and expressed as µmol βOHB turned over/min/kg-LBM. Because the liver is the primarily site of systemic ketone body production, whole-body turnover rates of ketone bodies at steady-state are mathematically equivalent to hepatic endogenous βOHB production rates **(V_RaβOHB_).**

### Quantifying Whole-body Glucose Turnover

Glucose turnover was calculated at metabolic and isotopic-steady state from the dilution of intravenously infused [3,4-^13^C_2_]glucose. The fractional enrichment of plasma glucose was determined from the ^13^C NMR spectra of MAG, by measuring the ratio (R) of enrichment at C3 and C4, relative to the natural abundance of the methyl groups from the acetone moiety of the MAG derivative. Fractional ^13^C enrichment (L_p_) of the plasma glucose pool was then calculated using linear regression, from a standard curve of known glucose enrichments and the equation L_p_ = 1.33 (R) + 0.031. Assuming that the fractional ^13^C enrichment of infused [3,4-^13^C_2_]glucose (L_i_) was 99%, and the known tracer infusion rate (F), whole body glucose turnover at metabolic and isotopic steady state was calculated as:

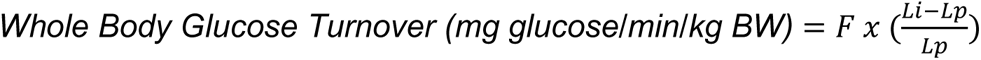

Using the DXA scan information, final flux units were expressed as µmoles glucose turned over/min/kg-LBM. In the fasted state the majority (>90%) of whole-body glucose production is derived from the liver, therefore total glucose turnover rates are mathematically equivalent to total hepatic EGP rates **(V_EGP_).**

### Quantifying Hepatic Glucose Sourcing Fluxes

Using the ^2^H enrichment patterns encoded in plasma glucose, the fractional contribution of various glucose sourcing pathways were calculated based on the following equations:

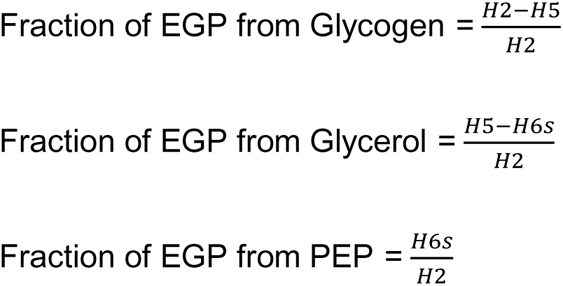

Where H2, H5 and H6s correspond to the relative ^2^H enrichment at the 2, 5 and 6s carbons of MAG. By multiplying each fraction by total EGP **(V_EGP_),** flux through each hepatic glucose sourcing pathway was calculated as follows:

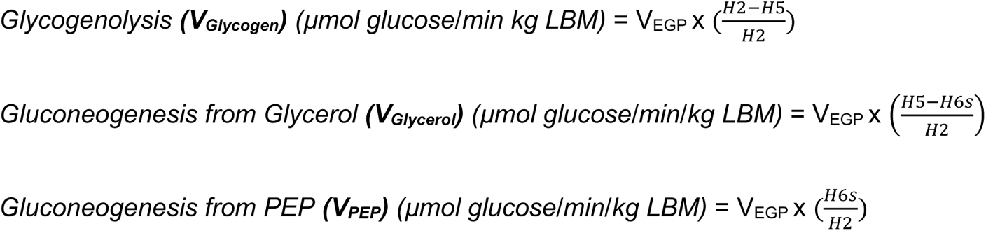

### Quantifying Hepatic Oxidative Fluxes

The rate of hepatic TCA cycle turnover, anaplerosis, cataplerosis and pyruvate cycling, can be modeled in the liver using the ^13^C isotopomers that arise in plasma glucose from orally administered [U-^13^C_3_]propionate. Briefly, at metabolic steady-state, glucose-linked rates of anaplerosis must equal cataplerosis, which are modeled relative to the turnover rate of the hepatic TCA cycle, assumed to be equal to the rate of CS (V_CS_). Because the non-oxidative entry (anaplerosis) and exit (cataplerosis) of carbon through the TCA cycle supports glucose anabolism via PEP-sourced GNG, changes in oxidative fluxes distort the ^13^C enrichment patterns of PEP which are transmitted and encoded in the multiplet resonances of plasma glucose. Using the known rate of PEP-derived GNG (V_PEP_), quantified as described above, and the ^13^C isotopomer populations quantified via ^13^C NMR, the absolute rate of the TCA cycle turnover (V_CS_), as well as anaplerotic (V_PC_) and cataplerotic fluxes (V_PEPCK_), and label recycling rates (V_PK+ME_) can be calculated as follows:

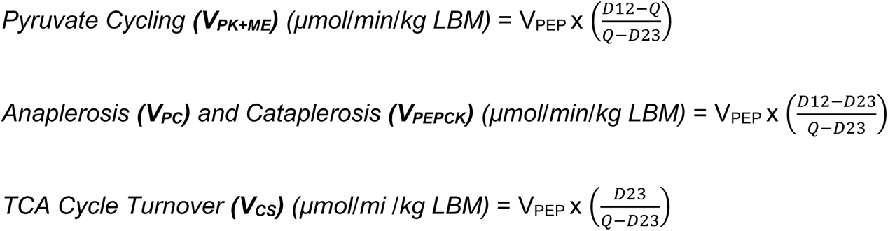

where D12 represents the doublet arising when C1 and C2 of glucose are labeled, D23 represents the doublet arising when C2 and C3 are labeled, and Q is the quartet arising when C1, C2 and C3 are all labeled. Pyruvate cycling (V_PK+ME_) reflects the summed rate of pyruvate kinase (PK) and malic enzyme (ME). Total anaplerosis is modeled as flux of PC (V_PC_), while cataplerosis is modeled as flux through PEP carboxykinase (PEPCK) (V_PEPCK_), which at steady-state are equal (V_PC_ = V_PEPCK_).

### Quantifying Hepatic Fat Oxidation Rates

By assuming that the liver is in a metabolic steady-state, the rate of acetyl-CoA production via β-oxidation can be assumed to be equal to the rate of acetyl-CoA disposal, thereby allowing total fat oxidation rates to be indirectly quantified as the sum of all disposal routes for hepatic acetyl-CoA. In the fasted liver, β-oxidation derived acetyl-CoA is (1) principally converted to ketone bodies which are largely released from the liver, (2) terminally oxidized in the TCA cycle, or (3) secreted as acetate. By quantifying each disposal pathway for acetyl-CoA, the majority of acetyl-CoA disposal can be accounted for, and used as a surrogate proxy of total acetyl-CoA turnover and fat oxidation at steady-state by the equation:

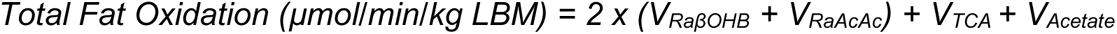

Fat oxidation is expressed in µmoles of acetyl-CoA turned over/min/kg-LBM. V_RaβOHB_ was measured via a tracer dilution of intravenously infused [U-^13^C_4_]βOHB. V_RaAcAc_ was calculated from V_RaβOHB_ turnover rates and the ratio of βOHB:AcAc. Acetate production rates (V_Acetate_) was assumed to be negligible in humans.

### Quantifying Hepatic RE Production

The production of RE’s, principally NADH and FADH_2,_ can be estimated from the above fluxes as previously described.^35,69^ This approach assumes that all acetyl-CoA originates from palmitate, and that the sourcing of V_PEP_ reflects 90% lactate and 10% pyruvate/alanine, in accordance with the typical hepatocellular redox state. The model assumes that complete β-oxidation of palmitate, a 16-carbon saturated FA, yields a total of 8 acetyl-CoA units, requiring 7 rounds of the β-oxidation cascade which collectively yields 14 REs, or an average of 1.75 REs per molecule of acetyl-CoA produced (14 REs / 8 Ac-CoA = 1.75 REs / Ac-CoA). By quantifying the rate for each disposal pathway for acetyl-CoA, the turnover of REs generated from fat oxidation can be calculated from the stoichiometry of each pathway as follows. For every mole of AcAc produced, two moles of acetyl-CoA turned over, and thus total REs generated from β-oxidation that are accounted for by AcAc **(RE_AcAc_)** = 3.5 x V_RaAcAc_. For every mole of βOHB produced, two moles of acetyl-CoA turned over, but conversion of AcAc to βOHB consumes 1 RE, and thus net REs generated from β-oxidation that are accounted for by βOHB **(RE_βOHB_)** = 2.5 x V_RaβOHB_. For every turn of the TCA cycle, one molecule of acetyl-CoA is consumed by CS, accounting for 1.75 REs, but each turn also generates an additional 4 REs, and thus total REs accounted for by the TCA cycle **(RE_TCA-total_)** = 5.75 x V_CS_. RE_TCA-total_ can be broken down into (1) total RE’s generated from β-oxidation **(RE_βox-CS_)** = 1.75 x V_CS_, and (2) total REs generated exclusively from the dehydrogenases of the TCA cycle (not β-oxidation) **(RE_TCA-only_)** = 4 x V_CS._ Total REs generated from β-oxidation can then be estimated as the sum of REs accounted for by AcAc, βOHB and the TCA cycle dehydrogenases as follows:

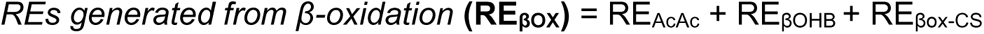

Total RE balance in the liver requires further accounting for GNG and three enzymes in particular: glycerol-3-phosphate dehydrogenase (G3PDH), glyceraldehyde-3-phosphate dehydrogenase (GAPDH), and lactate dehydrogenase (LDH). G3PDH generates 1 RE during GNG from glycerol. GAPDH consumes 1 RE during GNG from PEP, however, if the substrate used to source PEP is lactate, then LDH will generate 1 RE to offset this. Assuming that GNG is sourced 90% from lactate, and 10% from pyruvate + alanine, then total REs generated from GNG **(RE_GNG_)** = V_Glycerol_ – (0.1 x V_PEP_). From this, total REs turned over in the liver can then be calculated at steady-state as the sum of REs generated from β-oxidation, TCA cycle flux and GNG:

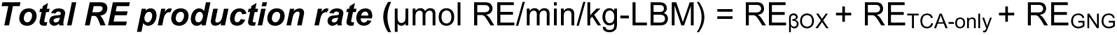

### Animal Models and Diets

All animal experiments were approved by the Institutional Animal Care and Use Committee at the University of Minnesota. All mice were bred on a C57BL/6NJ background, and received autoclaved water ad libitum. Mice were housed on corncob bedding in groups of 4 to 5 with lights off between 2000 and 0600 in a room maintained at 22°C. Sex of mice studied are indicated within each figure legend. Hepatocyte-specific HMGCS2 null mice were generated as described previously by crossing homozygous *Hmgcs2^flox/flox^* mice, with heterozygous Cre recombinase expressing mice, driven by the albumin promoter^70^. Littermate wild-type (WT) Alb-Cre-negative (*Hmgcs2^flox/flox^*, Alb-Cre^+/+^) animals were used as controls for Cre-positive (*Hmgcs2^flox/flox^*, Alb-Cre^+/Tg^) HMGCS2 KO mice, which are maintained on a C57BL6/NJ substrain hybrid background. *Hmgcs2* ASO-mediated loss of function was initiated in 6-week-old mice by i.p. injection (25 mg/kg) with murine *Hmgcs2*-targeted ASOs or scrambled sequence control ASOs (Ionis Pharmaceuticals) biweekly as described previously^42^. Hepatocyte-specific BDH1 null mice were generated as previously described^50^, by crossing homozygous *Bdh1^flox/flox^* mice^71^ to heterozygous Cre recombinase expressing mice, driven by the albumin promoter^70^. Littermate WT Cre-negative (*Bdh1^flox/flox^*, Cre^+/+^) animals were used as controls for Cre-positive (*Bdh1^flox/flox^*, Cre^+/Tg^) BDH1 KO mice. Food and water were provided ad libitum to mice in temperature-and humidity-controlled conditions maintained on a 14:10 h light/dark cycle. Mice were fed either a standard chow diet, or a high-fat diet. Standard chow diet (Teklad Global, 2016) consisted of 12% kcal from fat, 66% kcal from carbohydrates and 22% kcal from protein. Western diet (Envigo, TD.88137) consisted of 42% kcal from fat, 43% kcal from carbohydrates and 15% kcal from protein. 60% high-fat diet (Research Diets, D12492) consisted of 60% kcal from fat, 20% kcal from carbohydrates, and 20% kcal from protein. HFCR diet (Envigo, TD.160153.PWD) consisted of 90.5% kcal from fat, 0% kcal from carbohydrates and 9.5% kcal from protein.

### Quantifying Liver TAGs

20 mg frozen liver tissue was homogenized in 1 mL of 0.1x PBS, then total protein was quantified via BCA assay. Total lipids were extracted from 0.5 mg of protein via a Bligh-dyer lipid extraction, then total lipids were quantified colorimetrically using Infinity TAG reagent (Thermo, TR22421).

### Quantifying Liver Glycogen Content

Total glycogen was assayed as described previously.^49^ Briefly, 100 mg of frozen liver tissue was digested in 300 μL of 30% KOH (w/v), then glycogen was precipitated followed by enzymatic digestion using 0.3 mg/mL amyloglucosidase in 0.2 M sodium acetate buffer at pH 4.8 (Sigma 10115, 70 U/mg). Total glucose was quantified using ^1^H NMR then total glycogen was expressed in glucose units per mg of liver tissue.

### Serum NEFA, TAG and Blood Glucose Measurements

Serum NEFAs (Wako, 633-52001) and TAGs (Thermo, TR22421) were measure colorimetrically using commercial kits. Blood glucose was measured using glucose meters (Metene).

### Quantification of Total Ketone Bodies via UHPLC-MS/MS

Ketone bodies were formally quantified using UHPLC-MS/MS as described previously^50,72^. Briefly, [U-^13^C_4_]AcAc and [3,4,4,4-D_4_]βOHB internal standards were spiked into ice cold MeOH:ACN (1:1), then ketones were extracted from serum, perfusate, plasma, or sample extracts, followed by separation via reverse-phase UHPLC, and detected via parallel reaction monitoring (PRM) on a QExactive Plus hybrid quadrupole-orbitrap mass spectrometer. Quantification of ^13^C labeled ketone bodies was achieved post reduction with NaBD_4_ as previously described.^45,72^

### Quantification of Glucose and βOHB via ^1^H NMR

1.8 mL of liver effluent were collected, snap frozen and stored at -80°C till analysis. To prepare samples, the solvent was removed at 45°C under vacuum, then samples were reconstituted in 550 μl of ^2^H_2_O (99.9%) spiked with 0.3 mM d8-trimethyl-silyl propionate (TSP). ^1^H-NMR signals were acquired using a Bruker Avance III 600 MHz NMR instrument equipped with a CryoProbe, then the integrated intensities of the α-anomeric proton on glucose carbon-1, the methyl signal for βOHB, and the tri-methyl signal from TSP, were used to calculate molar concentrations of the respective substrates. For all ^1^H-NMR collections, spectra were collected by conventional pulse-and-collect measurements under quantitative conditions (10-ppm spectral range using ∼15 μs [90°] excitation pulse and 22-second delay between each of 20 transients).

### Stable Isotope Infusions in Mice

Jugular vein and carotid artery catheters were implanted into mice for isotope infusion and sampling as previously described.^56,73,74^ The exteriorized ends of the implanted catheters were flushed with 200 U·mL^−1^ heparinized saline and sealed with stainless-steel plugs. After surgery, mice were housed individually for ∼7 days of postoperative recovery before stable isotope infusions studies. All mice were within 10% of presurgical weight before stable isotope infusions. Mice were fasted during the duration of the infusions, and mice were fasted a total of 7 hours when samples were collected. Two hours into the fast, the exteriorized catheters were connection to infusion syringes. After a 1-hour acclimation period, an arterial blood sample (80 µL) was collected to determine glucose natural abundance enrichment, followed by a stable isotope infusion protocol to measure glucose-linked hepatic oxidative fluxes as previously described.^55,56^ Briefly, a ^2^H_2_O (99.9%)-saline bolus containing [6,6-^2^H_2_] glucose (99%) was infused over 25 min to enrich body water and prime (440 μmol·kg^−1^) the glucose pool. This was followed by a continuous infusion of [6,6-^2^H_2_] glucose (4.4 μmol·kg^−1^·min^−1^) till the end of the experiment. A primed (1.1 mmol·kg^−1^), continuous (0.055 mmol·kg^−1^·min^−1^) intravenous infusion of [U-^13^C] propionate was started 2 h after the ^2^H_2_O bolus. 100 µL arterial samples were collected at 90, 100, 110, and 120 mins after the [U-^13^C_3_] propionate bolus to determine arterial glucose concentration and enrichment. Donor erythrocytes were provided by constant rate infusion throughout the experiment to prevent a decline in hematocrit. Mice were euthanized by cervical dislocation following the final sample collection, then tissues were rapidly excised, freeze-clamped in liquid nitrogen. All samples were stored at −80°C.

### Glucose Derivatization, GC-MS Analysis and in vivo ^2^H/^13^C Metabolic Flux Analysis in conscious unrestrained Mice

Approximately 40 µL of plasma at each time point was used to synthesize di-*O*-isopropylidene propionate, aldonitrile pentapropionate, and methyloxime pentapropionate derivatives of glucose, then GC-MS analysis was utilized to collect uncorrected mass isotopomer distributions (MIDs) for six fragment ions as previously described.^54,55,75^ The metabolic flux analysis methodology used in these studies followed that previously described with minor modifications.^54^ To summarize, a metabolic reaction network was constructed using Isotopomer Network Compartmental Analysis (INCA) software.^76^ The reaction network defined the carbon and hydrogen atom transitions for hepatic glucose and associated oxidative reactions. The reaction velocities (i.e., flux) through each network reaction was determined relative to CS flux (V_CS_) by minimizing the sum of squared residuals between simulated and experimentally determined MIDs of the six fragment ions described above. Flux estimates were repeated 50 times from random initial values. Goodness of fit was assessed by a Chi-square test (*P* = 0.05) and confidence intervals of 95% were determined as previously described.^56^ Mouse body weights and the [6,6-^2^H_2_]glucose infusion rate were used to determine absolute values. In the GCMS-based model, total EGP (V_EGP_) is equal to the sum of glycogenolysis (V_PYGL_) and GNG (V_Aldo_), which are modeled as the rates of glycogen phosphorylase (PYGL) and aldolase (Aldo), respectively. Sourcing of GNG from glycerol (V_GK_) and PEP (V_Enol_) are modeled as the rates of glycerol kinase (GK) and enolase (Enol), respectively. Pyruvate cycling (V_PK+ME_) is equal to the summed rate of PK and ME and total cataplerosis is still equal to flux through PEPCK (V_PCK_). Anaplerosis is equal to the sum of PC (V_PC_) and propionyl-CoA carboxylase (PCC) (V_PCC_), and accounts for entry of unlabeled substrates through LDH (V_LDH_). Turnover of the 1^st^ span of the TCA cycle is modeled as the rate of CS (V_CS_), whereas turnover of the 2^nd^ span of the TCA turnover is modeled as the rate of succinate dehydrogenase (SDH) (V_SDH_), and reflects the sum of V_CS_ and V_PCC_.

### Mouse Liver Histology

Following euthanasia, mouse liver sections were fixed in 10% neutral buffered formalin, then tissue embedding, sectioning, and staining with hematoxylin and eosin was performed by the University of Minnesota Clinical and Translational Science Institute histology core.

### Quantifying Energy Charge, Redox State and Short-chain Acyl-CoAs

Energy nucleotides (ATP, ADP and AMP), redox nucleotides (NAD^+^ and NADH) and high-energy thioester containing acyl-CoAs (acetyl-CoA, propionyl-CoA and succinyl-CoA) were measured as previously described^50,77^, with modifications. Briefly, metabolites were extracted, separated using ion-pairing reverse-phase UHPLC-MS/MS on a C_18_ column (Waters Xbridge, 150 x 2.1mm, 3µm), then detected on a Thermo QExactive Plus mass spectrometer, operated in positive ionization mode, using PRM transitions as previously described^50,77^. Relative pool size (ion counts) normalized to biomass (mg total protein) was quantified for all analytes. Nucleotides were detected as adducts of dibutylamine acetate. Energy charge and Redox State was then estimated in the liver using the pool size of nucleotides and the equations:

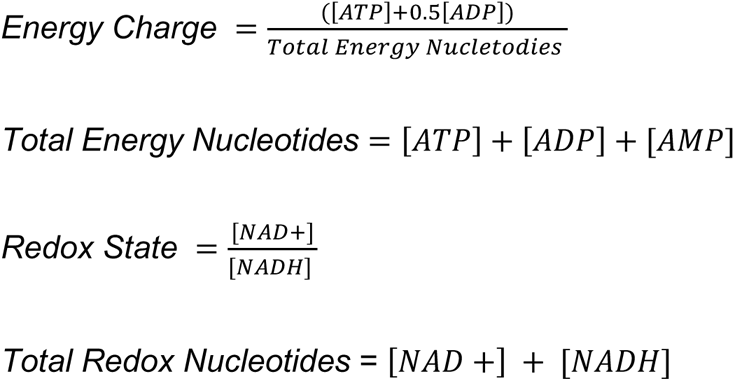

### Ex vivo Mouse Liver Perfusions

Liver perfusions were performed as described previously^49^. Briefly, mice were anesthetized with i.p. sodium pentobarbital injection, externally sterilized with 70% ethanol, then mice were dissected to expose the hepatic portal vein, followed by cannulation with a 24-gauge catheter needle. The cannula was secured, then the abdominal aorta, inferior vena cava and cardiac right atrium were severed to isolate the liver from peripheral tissues and prevent recirculation. Livers were perfused with oxygenated Krebs-Henseleit buffer (118 mM NaCl, 4.7 mM KCl, 2.5 mM CaCl_2_, 1.2 mM KH_2_PO_4_, 1.22 mM MgSO_4_, 25 mM NaHCO_3_, pH 7.4), warmed to ∼45°C and supplemented with 1.5 mM sodium lactate, 0.15 mM sodium pyruvate, 0.25 mM glycerol, 0.1 mM [U-^13^C_3_] propionate and 3% ^2^H_2_O. Perfusion buffers were also supplemented with exogenous FAs. Either 0.2 mM sodium octanoate was added directly to buffers, or 0.8 mM of a LCFA mixture (30% palmitate, 35%oleate, 8% stearate, and 27% linoleate) was bound to FA-free BSA at a BSA:FA molar ratio of 1:3, then dialyzed using a capillary dialysis filtration system and a Reveclear-300 dialysis cartridge (Baxter, 114745L) prior to addition to buffer. Specifics of buffer composition can be found in each figure legend. Perfusion buffer was oxygenated by continuous bubbling of a 95% O_2_, 5% CO_2_ gas mixture through the buffer reservoir using a fritted glass tube, and was delivered using a peristaltic pump at a flow rate of ∼8 mL/min for 60 minutes. Buffer and effluent samples were collected every 10 minutes. At the completion of the perfusion, the liver was freeze-clamped in liquid nitrogen, and subsequently weighed to obtain total liver weight for flux normalizations. Perfusion buffer (input), effluent (output) and tissues were stored at -80°C until analysis.

### Flux Calculations in Perfused Livers

Stable-isotope based flux modeling used *in vivo* in humans and rodents as described above was used to study metabolism of the *ex vivo* perfused liver within minor modifications as described previously.^49^ Briefly, total EGP rates **(V_EGP_)** was calculated from the rate of appearance of unlabeled glucose in the perfusate. βOHB production **(V_RaβOHB_)** and AcAc production **(V_RaAcAc_)** was calculated from the rate of appearance of unlabeled ketones in the perfusate. Livers were perfused with ^2^H_2_O and [U-^13^C_3_]propionate, which generate positional isotopomer populations that allow intrahepatic fluxes to be quantified as described during *in vivo* experiments in humans and mice. The ^2^H-enrichment of glucose at H2, H5 and H6s reports on the fractional contribution of glycogen, glycerol and PEP to glucose production, as described above. V_EGP_ measured from total glucose production in the perfusate, therein allows V_Glycogen_, V_Glycerol_ and V_PEP_ to be quantified as described above. The ^13^C-enrichment of glucose at carbons C1, C2 and C3 from [U-^13^C_3_]propionate distort the multiplet nuclear magnetic resonances of the C2 carbon in a manner that quantitatively encodes the ratio of TCA cycle turnover relative to anaplerosis, cataplerosis and pyruvate cycling. Because at steady-state the oxidative metabolic network herein described is coupled to PEP-linked GNG, by normalizing fluxes to V_PEP_ (measured as described above), absolute fluxes through the TCA cycle, anaplerosis, cataplerosis and pyruvate cycling can be calculated.

### Hepatic Fat Oxidation Calculations in ex vivo perfused livers

In perfusions, total fat oxidation was calculated as:

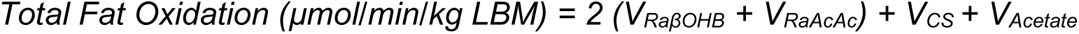

and expressed as μmol acetyl-CoA turned over/min/g liver. V_RaβOHB_, V_RaAcAc_ and V_CS_ are calculated as described above. Here, V_Acetate_ is quantified as the appearance of unlabeled acetate in the perfusate, and included in the *ex vivo LCFA* oxidation measurements. For octanoate perfused livers, it was assumed that very little acetate arose from octanoate, and therefore was not included in these calculations.

### RE Production Calculations in Liver Perfusions

The same theory and assumption used to model RE production rate *in vivo* in humans was used to model RE production rate in *ex vivo* perfused liver, with minor modifications. For LCFA perfusions, REs from each pathway were calculated using the same equations as above. For octanoate perfused livers, the following modifications were made. Because octanoate only generates 4 acetyl-CoA molecules, via three rounds of β-oxidation which generate 6 REs, the ratio of REs generated per acetyl-CoA molecule is 1.5 (6 RE / 4 Ac-CoA = 1.5), and not 1.75 as when palmitate is metabolized. Based on this the REs generated from each pathway can be calculated as follows:

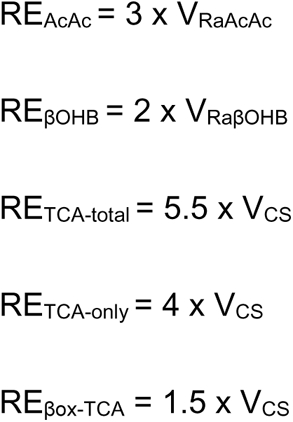

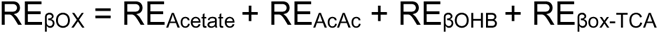

Total RE production rate can then be calculated based on V_Glycerol_ and V_PEP_ based on the assumptions that V_PEP_ is sourced 90% from lactate and 10% for pyruvate/alanine.

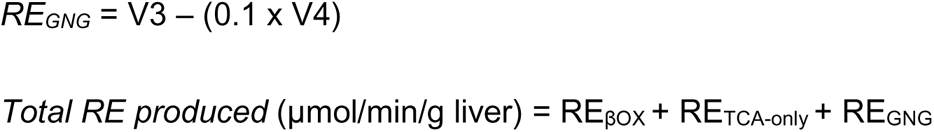

### Liver mitochondrial isolation

Mice were fasted for 18h with access to water. Livers were excised and placed in ice-cold mannitol-sucrose isolation media (MSIM; 220 mM Mannitol, 70 mM sucrose, 10 mM Tris, 1mM EDTA; pH=7.2) supplemented with 5 g/L fatty-acid free bovine serum albumin (BSA) (MSIM+BSA). Livers were minced in 5ml MSIM and liver pieces were rinsed three times in MSIM+BSA then transferred to an ice-chilled borosilicate glass vessel for homogenization. Homogenized liver tissue was centrifuged at 700 x g for 10 min at 4°C and the supernatant was filtered through two layers of gauze and centrifuged at 9,000 x g for 10 min at 4°C. The supernatant was discarded, and pellets were gently resuspended in 10 ml MSIM+BSA and centrifuged at 7,000 x g for 5 min at 4°C. The supernatant was discarded, and pellets were resuspended in 1.2 ml MSIM+BSA and centrifuged at 9,000 x g at 4°C. MSIM+BSA was aspirated from each tube and mitochondrial pellets were resuspended in 1.2 ml of MSIM without BSA and centrifuged at 9,000 x g at 4°C. MSIM without BSA was aspirated from each tube and mitochondrial pellets were resuspended in MSIM without BSA. Protein content was determined via a BCA protein assay and mitochondria were resuspended at a final concentration of 20 mg/ml for all functional assays.

### Mitochondrial respiratory control

Liver mitochondria were isolated as described above. Steady-state oxygen consumption rates (JO_2_) were determined using a modified version of the creatine kinase (CK) energetic clamp as described previously.^78^ Briefly, isolated mitochondria (0.150 mg/ml) were added to Buffer Z (105 mM MES potassium salt, 30 mM KCl, 10 mM KH_2_PO_4_, 5 mM MgCl_2_) supplemented with Creatine (5 mM), PCr (1.5 mM), CK (20 U/mL), fatty acid free BSA (2.5 mg/mL), and EGTA (1 mM) with either 10 μM Hymeglusin (HG) or the vehicle control (DMSO), followed by the addition of respiratory substrates (Pc/AKG; 50µM Palmitoyl-L-Carnitine, 5mM alpha-ketoglutarate), and ATP (5mM). Next, sequential additions of PCr was used to adjust the free energy of ATP hydrolysis (ΔG_ATP_). Following the respiration assays, 1.6mL of the mitochondrial buffer slurry was immediately collected and chilled on ice-water prior to differential centrifugation (9,0000g for 5 minutes at 4 °C) to separate the mitochondrial supernatant. Supernatants were flash frozen on dry-ice and stored at -80 °C until further analysis.

### Measuring mitochondrial NAD(P)H/NAD(P)+ redox

Fluorescent measurements of mitochondrial NAD(P)H/NAD(P)^+^ redox were determined using a QuantaMaster Spectrofluorometer (QM-400; Horiba Scientific) as described previously.^78^ All assays were conducted at 37°C in a 0.2ml reaction buffer. The buffer for all assays was Buffer Z supplemented with creatine (Cr; 5 mM), phosphocreatine (PCr; 1.5 mM), creatine kinase (CK; 20 U/ml), and either 10 μM Hymeglusin (HG) or DMSO (VCTRL). At the start of each assay, isolated liver mitochondria (0.4 mg/ml) were added to the assay buffer, followed by the addition of respiratory substrates (Pc/AKG) alone to capture 100% reduction within the NAD(P)H/NAD(P)^+^ couple, and then sequential additions of PCr to obtain the following final concentrations: 3, 6, 9, 12, 15 mM. After the final PCr addition, alamethicin (12.5ug/ml) was added to permeabilize the mitochondria. NAD(P)H/NAD(P)^+^ during the experiment was expressed as a percent reduced according to the formula: % Reduction = (F-F0%)/(F100%-F0%), where the 0% reduction state is represented as the fluorescent signal (Ex/Em, 340/450nm) recorded in the presence of alamethicin.

### Measuring Mitochondrial βOHB Production

Following the mitochondrial respiration control assays described above, 1.6mL of the mitochondria-buffer slurry was immediately collected and chilled on ice-water prior to differential centrifugation (9,000xg for 5 minutes at 4°C) to isolate the mitochondrial supernatant. βOHB levels in the supernatant were determined via a commercial kit (Wako Autokit 3-HB, Fujifilm Healthcare Solutions), according to manufacturer specifications, with modifications. Briefly, the principle of the assay is to colorimetrically monitor the reduction of NAD+◊NADH coupled to the oxidization of βOHB present in the assay samples. The reaction is catalyzed by the addition of βOHB dehydrogenase present in the R2 working reagent. To ensure AcAc does not interfere with the absorbance readings, AcAc in the samples is converted to acetone following the addition of acetoacetone decarboxylase present in the R1 working reagent. Prior to running the assay, mitochondrial supernatants were diluted 3-fold in Buffer Z. 20µL of mitochondrial supernatant samples and βOHB standard curve samples (0-12.5 µM) were loaded onto a 96-well flat-bottom plate in duplicate. 180µL of R1 working reagent was added to the standard and sample wells to monitor background absorbance at 405 nm for 5 minutes 37°C. Following, 180 µL/well of R2 working reagent was added, and absorbance was kinetically monitored 405 nm was monitored for 24 minutes at 37°C. For each of the βOHB standards, absorbance slope values (Δ absorbance/Δ seconds) were used to derive a linear standard curve to derive βOHB concentration present in each unknown sample. Unknown values were corrected for protein and total assay volume and expressed as nmol βOHB/mg protein.

### Mitochondrial incubations for metabolite profiling

Isolated liver mitochondria (0.02mg/mL) were incubated in 2mL Buffer Z supplemented with creatine (Cr; 5 mM), phosphocreatine (PCr; 1.5 mM), creatine kinase (CK; 20 U/ml), respiratory substrates (Pc/AKG), and either 10uM Hymeglusin (HG) or the vehicle control (VCTRL; DMSO), for 5 minutes at 37 °C with constant stirring. Subsequently, 5mM of ATP was added to reach respiratory state 3, followed by a 5-minute incubation at a fixed energy demand (ΔG_ATP_ = -14.26 kcal/mol) imposed by the addition of PCr (final concentration = 6mM). 1.6mL of the mitochondrial buffer slurry was immediately collected and chilled on ice-water prior to differential centrifugation (9,0000g for 5 minutes at 4 °C) to separate the mitochondrial pellet and supernatant. Both fractions were flash frozen on dry-ice and stored at -80 °C until further analysis.

### Mitochondrial acyl-CoA analysis by LC-MS/MS

Acyl-CoA analysis was performed as described in previous reports.^78–80^ Briefly, frozen mitochondrial pellets (0.4mg) were sonicated for ∼5 minutes in 1.1 ml extraction buffer (1:1 methanol/water containing 5% acetic acid) containing 20 μl 0.01 mM D9 pentanoyl-CoA as internal standard. Homogenates were centrifuged at 4°C for 15-minute at 18,000g and 900 μl of supernatant was used for solid phase extraction (SPE). For SPE, the cartridge (1-ml ion exchange cartridge packed with 100 mg of 2-2(pyridyl)ethyl silica gel (Sigma)) was pre-activated with 1 ml of methanol and then with 1 ml of extraction buffer (1:1 methanol/water containing 5% acetic acid). The acyl-CoAs trapped on the silica gel cartridge were released with (i) 1 ml of a 1:1 mixture of 50 mm ammonium formate, pH 6.3, and methanol and then (ii) 1 ml of a 1:3 mixture of 50 mm ammonium formate, pH 6.3 and (iii) 1 mL of methanol. The combined effluent was dried with nitrogen gas and stored at −80 °C until LC-MS analysis. A 40 μl sample was injected on a Pursuit XRs 5 C18 column (150 × 2.0 mm, 5μm), protected by a guard column (Pursuit XRs 5 C18 5 μm, 10 × 2.0 mm, 5μm) with temperature controlled at 25 °C, in an ExionLC AD liquid chromatograph system. The Analyst software (version 1.6) was used for data record and processing. Relative pool size estimates were determined by comparing analyte peak areas to D9 pentanoyl-CoA peak area, then normalized to the VCTRL (set to 1) to represent fold-change in the pool size of CoA in the presence of HG. The detailed LC-MS/MS conditions can refer to our previous reports.^78–80^

### Mitochondrial organic acid profiling by LC-MS/MS

Organic acids in the incubation buffer were methoximated and silylated to TBDMS-derivatives and profiled by a gas chromatography mass spectrometry (GCMS) method. Briefly, 200 μl of incubation buffer was spiked with 20 μl 0.05 mM norvaline as internal standard and then vortexed vigorously with 200 μl methanol and 1mL acetonitrile. Homogenates were then centrifuged at 4C for 15-minute at 18,000g and 675 μl of the resulting supernatant was dried under nitrogen gas at 37C. TBDMS-derivatives were prepared for GC-MS with MOX and TBDMS as described previously.^81,82^ GC-MS analyses were performed on an Agilent 7890B GC system equipped with a HP-5MS capillary column (30 m, 0.25 mm i.d., 0.25 µm-phase thickness; Agilent J&W Scientific), connected to an Agilent 5977A Mass Spectrometer operating under ionization by electron impact (EI) at 70 eV. Relative pool size estimates were determined by comparing analyte peak areas to norvaline peak area (m/z 288).

### Quantification of hydroxyalkenal species in liver tissue

Frozen liver tissue was homogenized in ice-cold water using 2.8-and 1.4-mm zirconium oxide beads at 4°C, then an appropriate amount of internal standard was spike in based on total protein determined using the BCA assay, followed by lipid extraction using a modified Bligh-dyer as previously described.^45,49^ Lipid extracts were then derivatized with carnosine and quantified using shotgun lipidomics as previously described.^83^

### Gene Expression

For mRNA expression analysis, samples were harvested and homogenized in Qiazol, then total RNA was isolated using Qiagen’s RNeasy protocol. First-strand cDNA was synthesized using iScript (BioRad, 170-8891), then quantitative real-time PCR was performed on a BioRad CFX384 Real-Time thermocycler using SsoAdvanced Universal SYBR Green supermix (BioRad, 170-5274). mRNA expression was calculated using the 2^-ΔΔCt^ method, where ΔCt = housekeeping gene Ct – gene of interest CT, then expressed as fold-change relative to the intact liver. mRNA levels were normalized to the intact liver, which was set to 1. Housekeeping gene was *Rpl32* for all targets. Primer sequences are given in **Supplemental Table 2.**

### Statistics and Data Analysis

All analyses were performed using GraphPad Prism version 9. Unpaired 2-tailed Student’s t-tests or 1-way ANOVA, was used to determine statistically significant differences. P-value < 0.05 was accepted as significant for all tests. Information about statistical analysis is included in each figure legend. Data are presented as mean ± SD, unless otherwise indicated.

## Acknowledgements

The authors acknowledge support from the National Institutes of Health (grants DK091538, DK122832, HL166142, DK136772, and AG069781). The authors are also incredibly grateful to the human participants who contributed to this study. The authors thank Rebecca Hollister, Nicholas G. Evanoff and the Center for Pediatric Obesity Medicine at the University of Minnesota Medical School for performing DXA scans. The authors also thank Oyedele Adeyi and Khalid Amin for liver histopathology reports.

## Acronyms

(HMG-CoA): 3-hydroxy-3-methylglutaryl-CoA
(HMGCS2): 3-hydroxy-3-methylglutaryl-CoA synthase 2
(AcAc): acetoacetate
(AIRg): acute insulin response to glucose
(ASO): antisense oligonucleotide
(BMI): body mass index
(BW): body weight
(BSA): bovine serum albumin
(CS): citrate synthase
(CRU): clinical research unit
(D-βOHB): D-beta-hydroxybutyrate
(DNL): de novo lipogenesis
(DXA): dual-energy X-ray absorptiometry
(EGP): endogenous glucose production
(FA): fatty acid
(FADH_2_): flavin adenine dinucleotide
(FFAs): free fatty acids
(FSIVGTT): frequently sampled intravenous glucose tolerance test
(GNG): gluconeogenesis
(GAPDH): glyceraldehyde-3-phosphate dehydrogenase
(G3PDH): glycerol-3-phosphate dehydrogenase
(HFCR) diet: high-fat carbohydrate-restricted
(HOMA-IR): homeostatic model assessment for insulin resistance
(HG): hymeglusin
(INCA): Isotopomer Network Compartmental Analysis
(KO): knock out
(LDH): lactate dehydrogenase
(LBM): lean body mass
(LCFAs): long chain fatty acids
(MRI): magnetic resonance imaging
(ME): malic enzyme
(MIDs): mass isotopomer distributions
(MS): mass spectrometry
(MASH): metabolic-dysfunction-associated steatohepatitis
(MASLD): metabolic-dysfunction-associated steatotic liver disease
(MAG): monoacetone glucose
(NAS): NAFLD activity score
(NADH): nicotinamide adenine dinucleotide
(NEFAs): non esterified fatty acids
(NAFLD): nonalcoholic fatty liver disease
(NMR): nuclear magnetic resonance
(PRM): spectroscopy parallel reaction monitoring
(PEP): phosphoenolpyruvate
(PUFAs): polyunsaturated fatty acids
(PDFF): proton density fat fraction
(PK): pyruvate kinase
(RE’s): reducing equivalents
(JO2): respiratory flux
(S/N): signal-to-noise
(SGLT2): sodium-glucose cotransporter 2
(MS/MS): tandem mass spectrometry
(TKBs): total ketone bodies
(TAG): triacylglycerol
(TCA) cycle: tricarboxylic acid
(FDA): U.S. Food and Drug Administration
(UHPLC): ultra-high performance liquid chromatography
(VLDL): very low-density lipoprotein

