## Supplemental material for "Ketogenesis protects against MASLD-MASH progression through mechanisms that extend beyond overall fat oxidation rate"

**Supplemental Figure 1. Stable isotope delivery protocol for *in vivo* flux modeling in bariatric MASLD-MASH patients.** For quantification of absolute flux through hepatic intermediary metabolic pathways subjects orally ingest three 250 mL doses of 70% heavy water ($^2H_2O$, $D_2O$) starting the night before flux measurements, then orally ingested two 300 mg doses of [U-$^{13}C_3$] propionate the morning of experiments. [3,4-$^{13}C_2$]glucose and [U-$^{13}C_4$]βOHB were then infused using a primed 2 hour continuous infusion with sample collection during the final 10 minutes after metabolic and isotopic steady state had been reached.

**Supplemental Figure 2. Expected relationships among metabolic fluxes in humans illustrate robustness of the model.** To demonstrate that metabolic fluxes followed logical biochemical patterns, Pearson correlation coefficients (r) were calculated to illustrate connections within the metabolic network. **(A)** Correlation matrix for the eight *in vivo* fluxes shown in heat map format, with the magnitude of correlation shown by color and r displayed in each box. As expected, **(B)** pyruvate cycling ($V_{PK+ME}$), total anaplerosis and cataplerosis ($V_{PEPCK}$) and TCA cycle turnover ($V_{CS}$) were all positively correlated, and **(C)** positively correlated with endogenous glucose production ($V_{EGP}$). **(D)** $V_{PK+ME}$, $V_{PEPCK}$, and $V_{CS}$ were positively correlated with the absolute flux of phosphoenolpyruvate (PEP) to glucose ($V_{PEP}$), but **(E)** were negatively correlated with fractional sourcing of glucose production from glycerol. Total fat oxidation was **(F)** strongly correlated with total ketogenesis ($V_{RaTKB}$), but **(G)** was not correlated with TCA cycle turnover ($V_{CS}$). Pearson correlations coefficients (r) are shown on each group along with a line of best fit and 95% confidence intervals calculated using linear regression. Correlations were accepted as significant if $p<0.05$ and are shown on each graph, or in each box on the correlation matrix an indicated, * $p<0.05$, ** $p<0.01$, ***$p<0.001$.

**Supplemental Figure 3. Endogenous ketogenesis correlates directly with liver injury in humans.** Correlation of NAS with absolute rates of hepatic glucose sourcing fluxes including **(A)** glycogenolysis ($V_{Glycogen}$), gluconeogenesis (GNG) from glycerol ($V_{Glycerol}$), and GNG from phosphoenolpyruvate (PEP) ($V_{PEP}$). **(B)** Correlation of NAS with absolute rates of pyruvate cycling ($V_{PK+ME}$) and total anaplerosis and cataplerosis ($V_{PEPCK}$). **(C)** Correlation of endogenous AcAc production ($V_{RaAcAc}$) and total ketone body production ($V_{RaTKB}$) with NAS score. $V_{RaAcAc}$ was calculated as the product of $V_{Ra\beta OHB}$ and the ratio of total AcAc : total βOHB. $V_{RaTKB}$ was calculated as the sum of $V_{RaAcAc}$ + $V_{Ra\beta OHB}$. **(D)** Correlation of NAS with the ratio of blood βOHB : AcAc. Correlation of $V_{Ra\beta OHB}$ with **(E)** PDFF**, (F)** AIRg, **(G)** serum AST, and **(H)** serum ALT. Pearson correlations coefficients (r) are given on each graph along with a line of best fit calculated using simple linear regression. Pearson correlations coefficients (r) are shown on each group along with a line of best fit and 95% confidence intervals calculated using linear regression. Correlations were accepted as significant if p<0.05 and are shown on each graph. For the correlation matrix, Pearson correlations coefficients (r) are shown in heat map format and are written in each square. If the p-value was significant it is displayed in the box as *. * p<0.05, ** p<0.01, ***p<0.001, as indicated.

**Supplemental Figure 4. Loss of hepatocyte HMGCS2 diminishes total liver adenylate nucleotides in HFCR diet-fed mice.** Quantification of liver energy and redox adenylate nucleotides in WT and hepatocyte-HMGCS2 KO mice fed HFCR diet for ~1 week. **(A)** Pool size of individual energy adenylate nucleotides (ATP, ADP, AMP), and **(B)** total pool of all energy adenylate nucleotides calculated as [ATP] + [ADP] + [AMP] (n=4-8/group). **(C)** Liver energy charge calculated as ([ATP]+0.5[ADP]) / ([ATP]+[ADP]+[AMP]) (n=4-8/group). **(D)** Pool size of individual redox adenylate nucleotides ($NAD^+$ and NADH), and **(E)** total pool of all redox adenylate nucleotides calculated as [$NAD^+$] + [NADH] (n=4-8/group). **(F)** Liver redox state calculated as [$NAD^+$] / [NADH] (n=4-8/group). Data are expressed as mean ± standard deviation. Statistical differences were determined by Student's t-tests and accepted as significant if p<0.05. * p<0.05, ** p<0.01, ***p<0.001, as indicated. NS = not statistically significant.

**Supplemental Figure 5. Loss of hepatocyte BDH1 does not alter body weight, liver weight or fasting blood glucose in western diet-fed mice.** Littermate control (WT) and BDH1-Liver-KO mice were fed a 42% high-fat western diet for ~16 weeks. **(A)** 18hr fasted serum ketone body analysis, and **(B)** longitudinal random-fed weight gain (n=11-12/group). **(C)** 18hr fasted liver weight and **(D)** blood glucose. Data are expressed as mean ± standard deviation. Statistical differences were determined by Student's t-tests and accepted as significant if $p<0.05$. * $p<0.05$, ** $p<0.01$, *** $p<0.001$, as indicated. NS = not statistically significant.

**Supplemental Figure 6. Loss of hepatocyte BDH1 does not worsen liver injury in high fat diet-fed mice.** Littermate control (WT) and BDH1-Liver-KO mice were fed a 60% high-fat diet (HFD) then **(A)** liver H and E (left) and picrosirius (right) histological staining was used to assess liver health. **(B)** Liver fibrosis was quantified from the picrosirius staining (n=4-5/group). **(C)** Expression of liver genes related to fibrosis and inflammation (n=4-5/group). Data are expressed as mean ± standard deviation. Statistical differences were determined by Student's t-tests and accepted as significant if $p<0.05$. * $p<0.05$, ** $p<0.01$, *** $p<0.001$, as indicated. NS = not statistically significant.

##### Table 1. MASLD-MASH Patient Characteristics

Table summarizing the human cohort studied encompassing markers of body composition, systemic glucose homeostasis and liver health.

##### Supplementary Table 1: Inclusion and Exclusion Criteria for Human MASLD-MASH Subjects

##### Supplementary Table 2: RT-qPCR Primer Sequences

Sequences of primers for RT-qPCR analysis of mRNA transcript abundance, given in the 5' to 3' direction.

### Supplemental Figure 1.

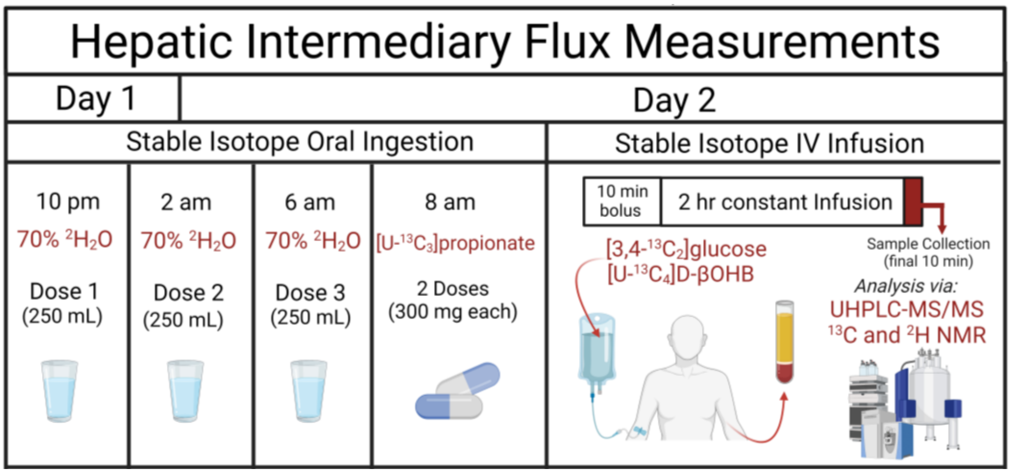

### Supplemental Figure 2.

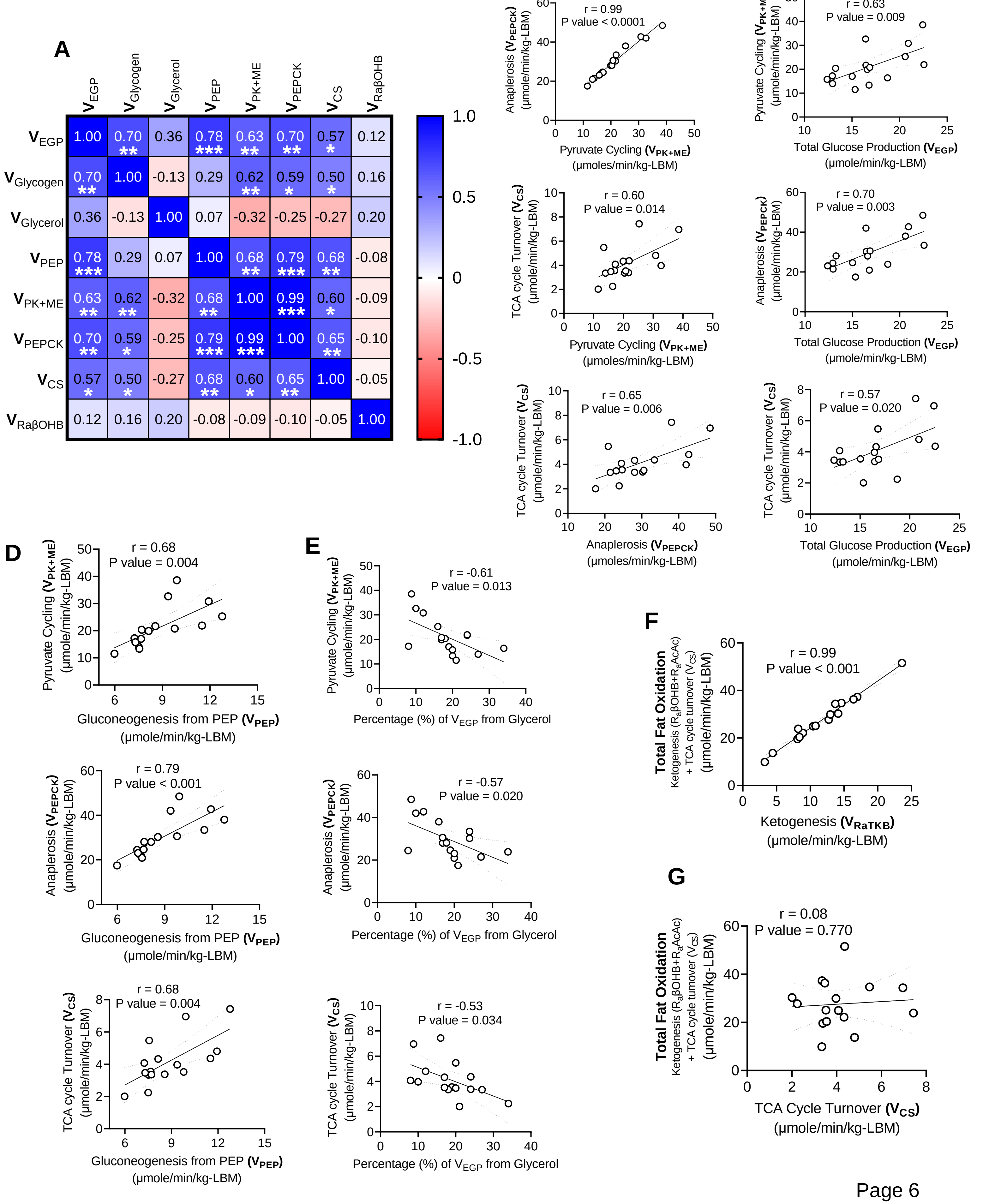

### Supplemental Figure 3.

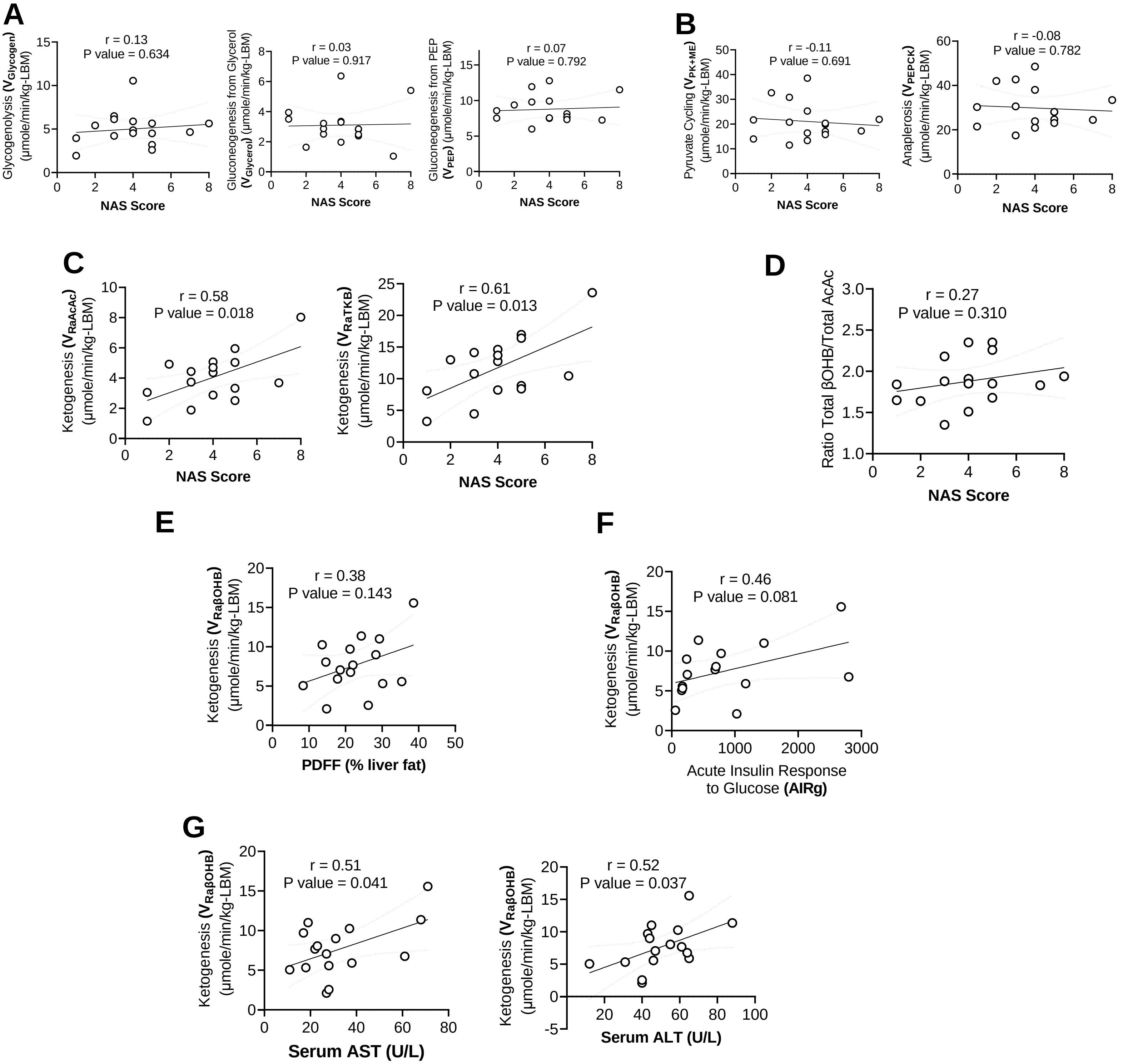

### Supplemental Figure 4.

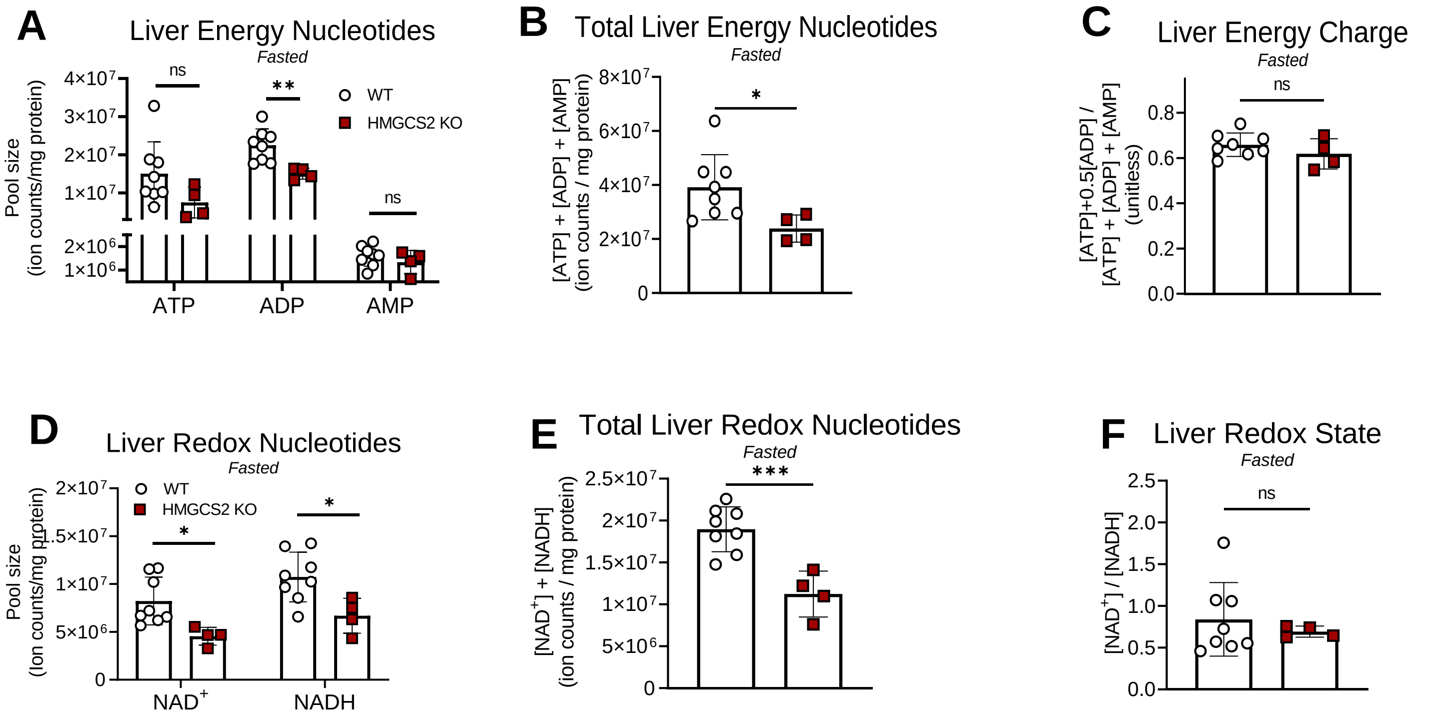

#### Supplemental Figure 5.

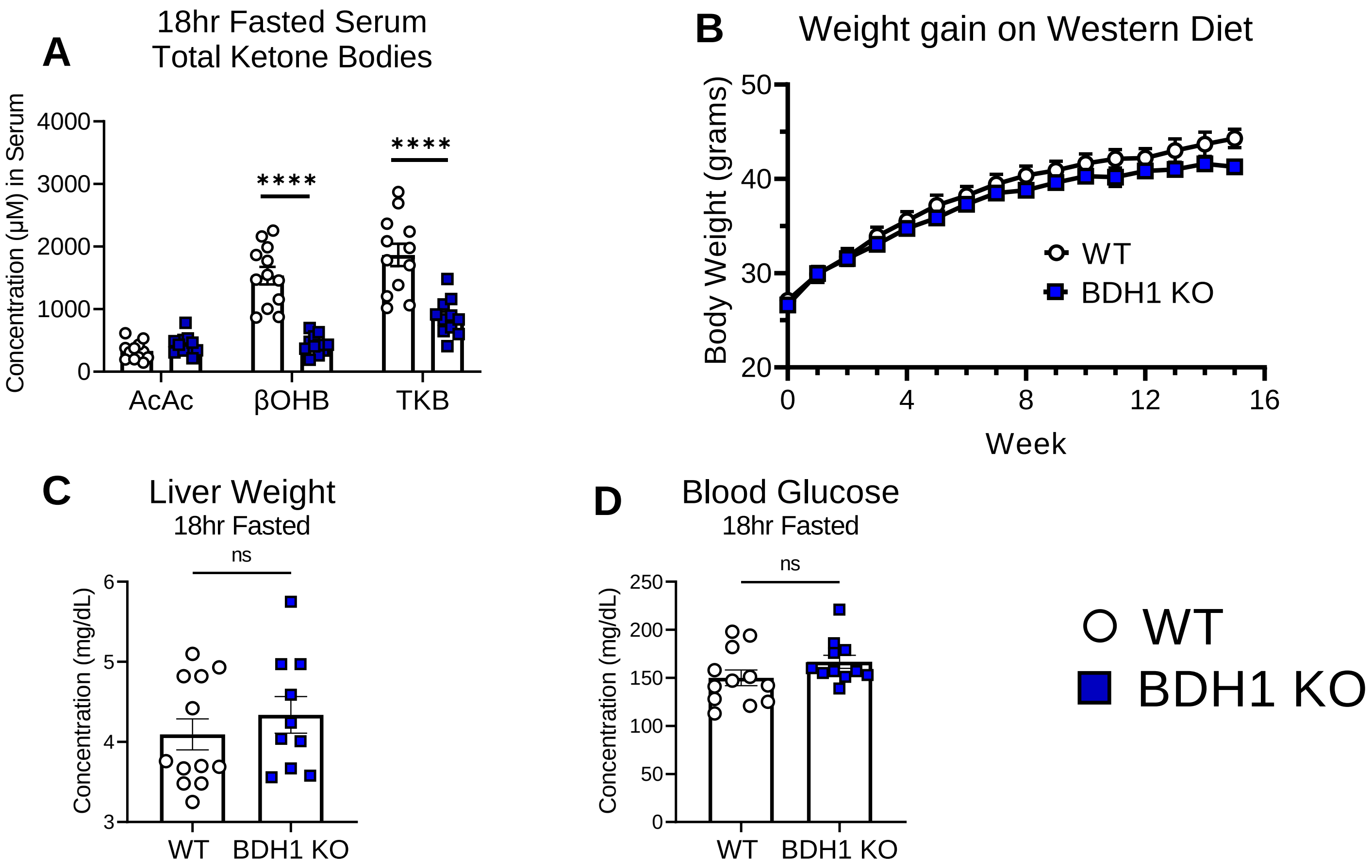

### Supplemental Figure 6.

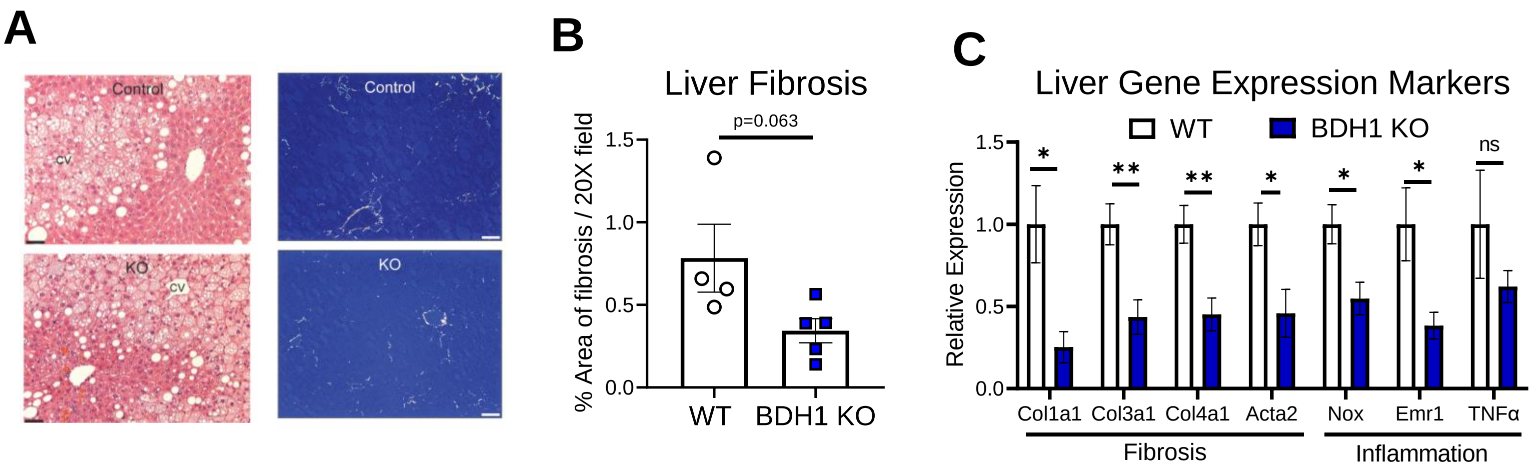

#### Table 1.

| Human Cohort Characteristics | | Average | ±SD |
|---|---|---|---|
| **Sex and Age** | Sex (F/M) | 15 / 1 | na |
| | Age (years old) | 48 | 14 |
| **Body Composition** | Body Weight (kg) | 112 | 18 |
| | Body Mass Index (BMI) (kg/m$^2$) | 42 | 6 |
| | Body Fat (% from DXA) | 51 | 4 |
| | Android Fat (% from DXA) | 59 | 4 |
| | Gynoid Fat (% from DXA) | 51 | 6 |
| | Lean Body Mass (LBM) (% from DXA) | 47 | 4 |
| | Bone Mineral Density (g/cm$^2$) | 1.3 | 0.2 |
| **Glucose Homeostasis** | Fasting Blood Glucose (mg/dL) | 104 | 13 |
| | Fasting Insulin (U/L) | 27 | 11 |
| | HOMA-IR | 6.7 | 2.6 |
| | HbA1c (%) | 5.8 | 0.5 |
| | Insulin Sensitivity (SI) ((mU/L)$^{-1}$ min$^{-1}$) | 3.8 | 8.1 |
| | Acute Insulin Response to Glucose (AIRg) ((mU/L) min$^{-1}$) | 851 | 843 |
| | Disposition Index (DI) (SI x AIRg) (unitless) | 1294 | 1056 |
| | Glucose Effectiveness ($S_g$) x10$^{-2}$ (min$^{-1}$) | 1.2 | 0.5 |
| | Glucose Effectiveness at Zero Insulin (GEZI) x10$^{-2}$ (min$^{-1}$) | 0.4 | 1.7 |
| | Beta-cell Function (mU/L) | 378 | 226 |
| | Insulin Resistance | 5.9 | 2.2 |
| **Liver Function** | Alanine Transaminase (ALT) (U/L) | 50.3 | 16.7 |
| | Aspartate Aminotransferase (AST) (U/L) | 32.9 | 17.7 |
| | Albumin (g/dL) | 3.9 | 0.3 |
| | Alkaline Phosphatase (ALP) (U/L) | 95.4 | 20.9 |
| | Bilibrubin (mg/dL) | 0.5 | 0.3 |

### Supplemental Table 1.

| |
|---|
| **1.1 Inclusion Criteria:** |
| 1. Age 30 to 67 years at eligibility visit |
| 2. At least one of the following: |
| a) Diagnosed with NASH with a total NAS ≥ 4 including a ballooning score of at least 1, or non-NASH/NAFLD with a total NAS <3 |
| b) Diagnosed with T2DM or prediabetes, HbA1c< 8% and NOT requiring insulin or other oral/injectable hypoglycemic agents |
| 3. Body Mass Index (BMI) 30.0-45.0 kg/m2 at eligibility visit |
| 4. Willingness to accept surgical intervention after an individual seminar session |
| 5. All patients must have insurance with no exclusion for obesity related treatments or management of obesity surgery complications. This applies to all patients enrolled in the study |
| 6. Expect to live or work within approximately one-hour traveling time from the study clinic for the duration of the one-year trial |
| 7. Willingness to comply with the follow-up protocol and successful completion of the run-in (described in section 5.2) |
| 8. Written informed consent |
| 9. Suitable for liver biopsy using the percutaneous approach |
| 10. Vulnerable populations will not be targeted for inclusion, but those noted in section 9.1 may be allowed to participate provided they met all of the inclusion and none of the exclusion criteria. |
| **1.1 Exclusion Criteria:** |
| 1. Cardiovascular event (myocardial infarction, acute coronary syndrome, coronary artery angioplasty or bypass, stroke) in the past six months. |
| 2. Current evidence of congestive heart failure, angina pectoris, or symptomatic peripheral vascular disease. |
| 3. Cardiac stress test indicating that surgery or IMM would not be safe. |
| 4. Pulmonary embolus or thrombophlebitis in the past six months. |
| 5. Cancer of any kind (except basal cell skin cancer or cancer in situ) unless documented to be disease-free for five years. |
| 6. Significant anemia (hemoglobin 1.0 g/dL or more below normal range) or history of coagulopathy. |
| 7. Serum creatinine >1.5 mg/dL. |
| 8. Serum total bilirubin greater than the upper limit of normal in the absence of Gilbert's syndrome, or alkaline phosphatase or ALT or AST greater than 2.5 the upper limit of normal. Elevated INR. |
| 9. Alcohol intake more than one drink or >20 grams per day |
| 10. History of stomach surgery, bile duct surgery, pancreatic surgery, splenectomy, or colon resection. |
| 11. Gastric or duodenal ulcer in the past six months. |
| 12. History of intra-abdominal sepsis (except for uncomplicated appendicitis or diverticulitis more than six months prior to enrollment). |
| 13. Previous organ transplantation. |
| 14. Self-reported HIV-positive status, active tuberculosis, active malaria, chronic hepatitis B or C, cirrhosis, or inflammatory bowel disease. |
| 15. Currently pregnant or nursing, or planning to become pregnant in the next two years. |
| 16. History of alcohol, drug, or opioid dependency (excluding nicotine) in the past five years. |
| 17. Active psychosocial or psychiatric problem that is likely to interfere with adherence to the protocol. |
| 18. Depression A CESD score more than 17 and a psychologist determination that the patient is not a good fit for surgery. |
| 19. Current participation in a conflicting research protocol. |
| 20. Presence of any chronic or debilitating disease that would make adherence to the protocol difficult. |
| 21. 12-lead EKG indicating that surgery would not be safe. |
| 22. Serum c-peptide <1.0 ng/ml post prandial. |
| 23. Exclusions may also be made at the discretion of the attending physician or the eligibility committee. |
| 24. Contraindication to MRI scanning. MRI contraindications are assessed by MR technologists on the day of scanning using a standard safety screening form. |
| 25. Gastroesophageal reflux disease requiring medications. History of endoscopy demonstrating esophagitis or Barretts changes in the esophagus. Any history of dysphagia. |
| 26. More than 2 cups of coffee per day. |
| 27. Treatment with drugs associated with nonalcoholic fatty liver disease (amiodarone, methotrexate, oral glucocorticoids at doses greater than 5 mg/day, tamoxifen, estrogens at doses greater than those used for hormone replacement or contraception, anabolic steroids, valproic acid) for more than 4 weeks within the last 2 months prior to the initial screening. |
| 28. Treatment with pioglitazone or high-dose vitamin E (>400 IU/day) within the last 2 months prior to the initial screening. |
| 29. Initiation of treatment with a glucagon-like peptide-1 (GLP-1) agonist or a dose change within the last 2 months prior to the initial screening |

### Supplemental Table 2.

| RT-qPCR Primer Sequences | | |
|---|---|---|
| **Gene** | **Forward Primer Sequence (5` --> 3`)** | **Reverse Primer Sequence (5` --> 3`)** |
| **Col1a1** | TGCTTCGTGTAAACTCCCTC | TTGTTCGTCTGTTTCCAGGG |
| **Col3a1** | GAAGTCTCTGAAGCTGATGGG | TTGCCTTGCGTGTTTGATATT |
| **Col4a1** | AGACCATTCAGATTCCGCAG | CGCTTCTAAACTCTTCCAGACAG |
| **Acta2** | CCGCAAATGCTTCTAAGTCC | TTTGGGCAGGAATGATTTGG |
| **Nox2** | AGAGCATCTCCAACTCAGAATCCG | ACACTGGTTTCCTGGTGAAAGAGC |
| **Emr1** | CTTTGGCTATGGGCTTCCAGTC | GCAAGGAGGACAGAGTTTATCGTG |
| **Tnfa** | TGGAGTCATTGCTCTGTGAAGGGA | GGGCTCTGAGGAGTAGACAATAAAGG |
| **Rpl32** | CCTCTGGTGAAGCCCAAGATC | TCTGGGTTTCCGCCAGTTT |
